# Fast, accurate parsing of antibody structures and antibody-antigen interactions enables a comprehensive structural antibody database

**DOI:** 10.1101/2025.02.25.640196

**Authors:** Xiaoqiang Huang, Jun Zhou, Shuang Chen, Xiaofeng Xia, Y. Eugene Chen, Jie Xu

## Abstract

Antibody (Ab) structures and antibody-antigen interactions (AAIs) are essential for understanding immune recognition and designing Ab therapeutics. While existing structural Ab databases provide valuable insights, they still have notable limitations. Here, we present SAAINT-parser, an computational workflow for rapid, accurate, and robust extraction of Ab and AAI information from the Protein Data Bank (PDB). SAAINT-parser features precise detection of Ab chains, accurate pairing of Ab chains, and reliable identification of AAIs. The resulting SAAINT-DB contains 18,031 data entries from 9,373 PDB structures, offering a comprehensive and up-to-date resource. Detailed analyses of SAAINT-DB reveals its advantages over the most widely used SAbDab database, particularly in terms of data accuracy and completeness. Furthermore, SAAINT-DB provides nearly twice as many nonredundant, manually curated Ab-Ag binding affinity data compared to SAbDab. Both SAAINT-parser and SAAINT-DB are available as open-source resources, aimed at advancing Ab-related research and benefiting the broader scientific community.

## Introduction

Antibodies (Abs), also known as immunoglobulins (Igs), are essential immune system components, providing defense against infections and foreign substances. Their high affinity and specificity for antigens (Ags) have made them invaluable in basic research, biotechnology, and biomedical applications^1,2.^ Structural biology and biophysical studies on Ab structures and Ab-Ag interactions (AAIs) are crucial for understanding the molecular determinants of Ab specificity, binding affinity, and other properties, facilitating the rational design and optimization of Abs for therapeutic and diagnostic purposes.

Recent breakthroughs in artificial intelligence, particularly deep learning, have revolutionized protein structure prediction and design. The availability of extensive, diverse protein structural data has played a pivotal role in these advancements. Trained on a vast portion of the Protein Data Bank (PDB)^3^, the Nobel Prize-winning AlphaFold^4^ system, related networks^5,6^ , and their successors^7,8^ significantly outperform traditional methods in modeling Ab structures and AAIs^9,10.^ Similarly, the RFdiffusion^11^ network, fine-tuned on Ab complex structures, has demonstrated the ability to accurately design de novo single-chain Abs^12^.

The PDB^3^ serves as the primary repository for 3D structural data of large biological molecules, including Abs. As of this writing in Feburary 2025, the PDB contains over 231,000 entries, yet only a small fraction (∼4%) contains Ab structures. To address this limitation, several specialized structural Ab databases^13–19^ have been developed, integrating data from the PDB and other resources. Among them, IMGT/3Dstructure-DB^13^ is one of the earliest databases dedicated to immunological proteins, including Abs. Other notable databases include BEID (*B*-cell *E*pitope *I*nteraction *D*atabase)^14^, which compiles sequence-structure-function data on AAIs, and AgAbDb (*A*nti*g*en-*A*nti*b*ody interaction *D*ata*b*ase)^16^, a knowledgebase developed to compile, curate, and analyze determinants of AAIs. PyIgClassify^17^ specializes in the classification of complementarity-determining regions (CDRs), while AB-Bind^18^ provides a set of binding mutational data with Ab structures for evaluating computational AAI modeling methods. AbDb (*A*nti*b*ody structure *D*ata*b*ase)^19^ focuses on the Ab Fv (*F*ragment *v*ariable) regions with their conjugate Ags.

Among these, SAbDab (*S*tructural *A*nti*b*ody *Da*ta*b*ase)^15^ stands out as the most comprehensive repository. According to its authors, SAbDab contains all the publicly available Ab structures annotated with key experimental details, heavy chain (HC)-light chain (LC) pairings, and Ag information, and manually curated Ab-Ag binding affinity data. As of its most recent update on January 17, 2025, SAbDab includes 9,200 PDB structures and 18,265 data entries, surpassing other structural Ab databases described above. In comparison, IMGT/3Dstructure-DB^13^, the second largest, last updated on May 23, 2024, contains 7,243 PDB structures and only 8,890 data entries. Given its breadth and regular updates, SAbDab currently represents the most extensive structural Ab database available.

While these databases provide valuable structural data for Ab-related research, they still have notable limitations. For instance, our detailed examination of SAbDab reveals a significant amount of missing or incorrectly annotated data, which can hinder accurate analysis and computational modeling.

To address these issues, we present SAAINT-parser, a computational workflow designed for fast and accurate processing of PDB entries to extract structural Ab and AAI information. This has enabled the construction of SAAINT-DB, a novel and comprehensive structural Ab database. Both SAAINT-parser and SAAINT-DB are released as open-source resources, designed to advance Ab-related research and benefit the broader scientific community.

## Results

### The SAAINT-parser workflow

The SAAINT-parser workflow extracts paired Abs, unpaired Ab chains, and AAIs from experimentally solved structures using their PDB identifiers (IDs) as input (Fig. 1a). It consists of three main modules that process the PDB-associated FASTA file, mmCIF file, and web content, respectively. These modules generate intermediate data, which are then integrated to infer the outcomes. Below is a description of the key components, with further details provided in the Methods section.

**Fig. 1.**
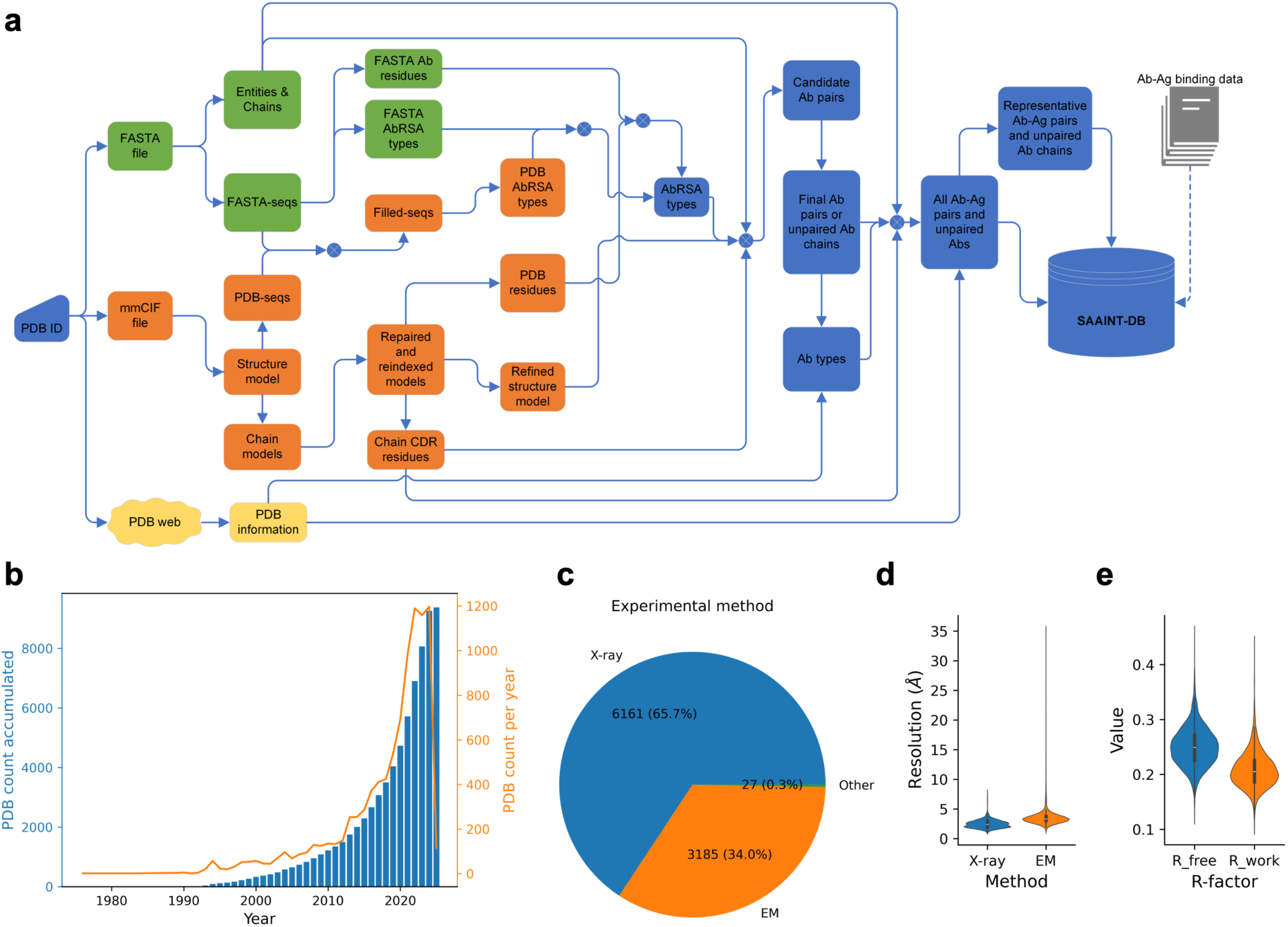
Overview of SAAINT-parser and SAAINT-DB. **a**, The SAAINT-parser workflow. **b**, Number of PDB entries in SAAINT-DB. **c**, Distribution of experimental methods for solving PDB structures in SAAINT-DB. **d**, Distribution of resolution for X-ray and electron microscopy (EM) structures. **e**, Distribution of R-factor values for X-ray structures.

The FASTA module processes chain entities and sequences from the FASTA file, utilizing AbRSA^20^ to determine whether a FASTA sequence (FASTA-seq) contains Ab variable domains and, if possible, to identify Ab residues (FASTA Ab residues). The mmCIF module employs Biopython^21^ to parse the complete structural model from the mmCIF file. Protein sequences (PDB-seq) consisting of residues with Cα atomic coordinates are extracted from the structure, aligned to the corresponding FASTA-seq, and used to generate a gap-filled protein sequence (Filled-seq), which then undergoes a second AbRSA analysis. The entire structure is further divided into individual chains, each reindexed based on the alignment for Filled-seq creation and repaired using Pulchra^22^ to restore missing atoms. Each repaired chain is analyzed to identify protein residues (PDB residues) with atomic coordinates and subjected to AbRSA_PDB^20^ to detect CDR residues if classified as an Ab chain. The repaired individual chains are then concatenated and optimized using FASPR^23^, which repacks protein side chains to minimize steric clashes. Additionally, the web module retrieves key PDB information by directly accessing the PDB webpage.

During the integration phase, AbRSA types inferred from FASTA-seqs and Filled-seqs are combined with data such as FASTA Ab residues and PDB residues to determine the final AbRSA types. Dependent on chain entities and final AbRSA types, all possible HC-LC pairs (HL pairs) are extracted from the FASPR model and analyzed with UniDesign^24,25^, yielding a set of relatively reliable HL pairs, which are further optimized by a combination of greedy search and iterative heuristic search to ensure that one HC is paired with at most one other LC or vice versa (see Methods). The final HL pairs or unpaired HCs/LCs, along with the web-fetched PDB title, are then used to determine Ab types.

The AAI identification process follows a similar approach to identifying HL pairs. For each Ab, all possible Ab-Ag pairs are extracted and analyzed by UniDesign. An Ab-Ag pair is considered valid if its interface contains a relatively higher number of residues (e.g., ≥10) and a higher proportion of CDR residues (e.g., ≥0.25). To ensure comprehensive representation, all Ag chains associated with a given Ab are merged.

All identified AAIs and/or Abs are recorded in the SAAINT-DB database. Since some structures contain multiple copies of the same Ab, a carefully designed scoring function is used to select the most representative AAI or Ab (see Methods). The collection of representative AAIs or Abs from various PDB entries is stored separately in SAAINT-DB.

Finally, besides the data generated by SAAINT-parser, manually curated Ab-Ag binding affinity data is also incorporated for representative AAIs based on relevant publications.

### The SAAINT-DB full dataset

The SAAINT-DB full dataset (SAAINT-DB-full) comprises 18,031 data entries derived from 9,373 PDB entries released between May 19, 1976 and January 29, 2025 (deposited between March 17, 1976 and December 20, 2024) (Fig. 1b). Over time, the number of experimentally determined Ab-involved PDB entries has steadily increased each year, with a notable increase between 2020 and 2023, likely driven by the global COVID-19 pandemic related research.

The dataset is predominantly composed of structures determined by X-ray diffraction (6,161 structures, 65.7%) and electron microscopy (EM) (3,185 structures, 34.0%), while other experimental methods account for only 27 structures (0.3%) (Fig. 1c). X-ray structures exhibit resolutions ranging from 0.92 to 8.0 Å with a median resolution of 2.4 Å (Fig. 1d). Among them, 3,539 (57.4%) have a moderate resolution of ≤2.5 Å, indicating moderate or better quality. The R-free and R-work values range from 0.122 to 0.458 (median: 0.249) and from 0.102 to 0.441 (median: 0.205), respectively (Fig. 1e). EM structures have resolutions spanning 1.7 to 35.0 Å, with a median of 3.3 Å (Fig. 1d). Of these, 2,920 (82.5%) have a resolution of ≤4.5 Å, indicating medium or better quality.

The basic structure of an Ab usually consists of HCs and LCs, each made up of Ig domains including VH (*V*ariable domain of *H*C) and VL (*V*ariable domain of *L*C), which contain the CDRs and FRs (*F*ramework *R*egions) and constant regions (CH and CL). Ab structure is functionally divided into two main regions: Fab (*F*ragment *a*ntigen-*b*inding) and Fc (*F*ragment *c*rystallizable). These regions are connected by a flexible hinge, allowing the Fab arms to move and bind to Ags at various angles. Fv, the smallest functional unit of an Ab responsible for Ab binding and specificity, consists of VH and VL.

The HCs and LCs of paired Abs may originate from different sources. Additionally, some Ab entries consist of a single chain, requiring separate source analysis for HCs and LCs. The top five species sources of HCs are *Homo sapiens* (8,744 entries), *Mus musculus* (4,373 entries), *Lama glama* (1,652 entries), *Vicugna pacos* (776 entries), and synthetic constructs (586 entries) (Fig. 2a). For LCs, the top sources are *Homo sapiens* (8,315 entries), *Mus musculus* (3,913 entries), *Macaca mulatta* (307 entries), *Oryctolagus cuniculus* (174 entries), and *Rattus norvegicus* (107 entries) (Fig. 2b).

**Fig. 2.**
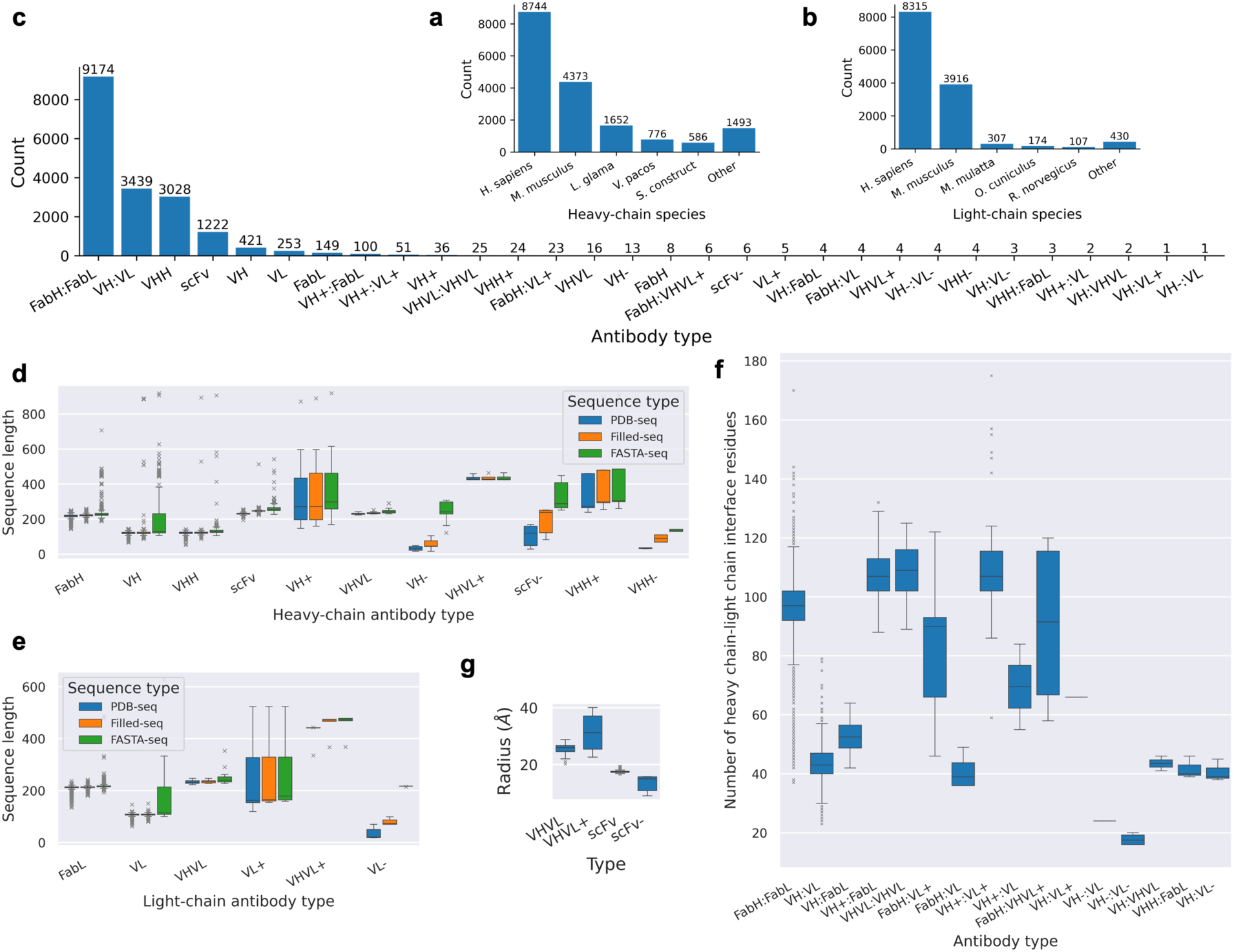
Statistics of antibodies in SAAINT-DB. **a**, Top sources of antibody heavy chains. **b**, Top sources of antibody light chains. **c**, Antibody type classification in SAAINT-DB. **d**, Distrition of sequence lengths for heavy-chain antibody types. **e**, Distribution of sequence lengths for light-chain antibody types. **f**, Distribution of the number of heavy chain-light chain interface residues across distinct antibody types. **g**, Distribution of the mean radius for scFv and VHVL types.

As with any structural Ab database, Ab structures form the foundation of SAAINT-DB. However, these structures, sourced from the PDB with minimal processing, lack a unified Ab format. Fully solved Abs, consisting of all Ab domains, are rare; instead, Fab and Fv are the most common formats. Another widely studied format is VHH (*V*ariable *H*eavy domain of *H*eavy chain only Ab), also known as a nanobody. Additionally, various engineered Ab formats exist, such as scFv (*s*ingle-*c*hain *Fv*). Due to experimental limitations, some Ab structures contain missing residues or domains, leading to discrepancies between deposited structures and their annotations. For example, an Ab is annotated as a Fab, whereas its atomic coordinates only include Fv domains. In this situation, it would be more appropriate to classify it as an Fv rather than a Fab.

To address this, SAAINT-parser implements a robust method for automatically categorizing Ab entries into the most appropriate types, such as Fab, Fv, VHH, scFv, and others, by considering sequence data, structural features, and PDB annotations (see Methods). To explicitly denote paired chains, we introduce chain-specific Ab types for both HCs and LCs. For instance, instead of simply labeling an Ab as Fab, we specify it as FabH:FabL. Similarly, we use VH:VL instead of Fv to represent paired variable domains. Overall, SAAINT-DB-full defines 30 Ab types; among them, the most common are FabH:FabL, VH:VL, VHH, scFv, VH, VL, FabL, and VH+:FabL, with respective counts of 9,174, 3,439, 3,028, 1,222, 421, 253, 149, and 100 entries (Fig. 2c). This aligns with the known prevalence of Fab, Fv, VHH, and scFv in solved Ab structures.

Protein chain length plays a critical role in classifying Ab types. However, determining Ab type solely from its FASTA-seq can be unreliable, as these sequences may include a large number of non-Ab residues. To improve classification accuracy, we analyze lengths from multiple sequence representations: FASTA-seq, PDB-seq, and Filled-seq, with a particular focus on the latter two. It is well established that VH, VHH, and VL domains are single-domain structures of approximately 110 amino acids (aa), whereas FabH and FabL include both a variable and constant domain, making them roughly twice as long (∼220 aa). As shown, many VH FASTA-seqs exceed 400 aa, whereas their PDB-seqs and Filled-seqs remain ∼110 aa (Fig. 2d). Similarly, while many FabH FASTA-seqs surpass 400 aa, their PDB-seqs and Filled-seqs remain ∼220 aa. Additionally, VH-, VL-, and scFv-(representing incomplete VH, VL, or scFv structures) are generally shorter than their parental types (Fig. 2d,e). In contrast, VH+, VHH+, VL+, and VHVL+ exhibit broader length distributions (Fig. 2d,e), as they usually represent chimeric constructs combining Ab domains with other protein regions.

For a given Ab type, the number of interface residues in HL pairs (*N_HL_inf_res_*) can vary significantly (Fig. 2f), reflecting differences in pairing angles^26^ and packing densities between HCs and LCs. For example, this number ranges from as few as 20 aa to as many as 80 aa in VH:VL types.

Both VHVL and scFv consist of a VH and a VL domain, and their PDB-seqs cluster ∼220 aa (Fig. 2d,e), making it difficult to distinguish them by sequences. However, a key structural difference is that scFv’s VH and VL domains pack more tightly, resulting in a smaller mean radius (see Methods). We found that a cutoff of 20 Å can effectively differentiate scFvs from VHVLs with perfect accuracy (Fig. 2g).

Among the 18,031 entries in SAAINT-DB-full, 13,663 (75.8%) are classified as AAIs, where each Ab interacts with one or more Ag chains. Of these, 10,990 (80.4%) involve a single Ag chain, 2,573 (18.8%) involve two, and 100 (0.8%) involve three, with no entries exceeding three Ag chains (Fig. 3a). The Ag chains include 12,204 proteins, 1,410 peptides, 16 DNAs, and 62 RNAs (Fig. 3b). Regarding Ag sources, the most common species are *Homo sapiens*, SARS-CoV-2, HIV-1, influenza A, and *Plasmodium falciparum*, contributing 5,032, 2,673, 1,559, 389, and 296 entries, respectively (Fig. 3c).

**Fig. 3.**
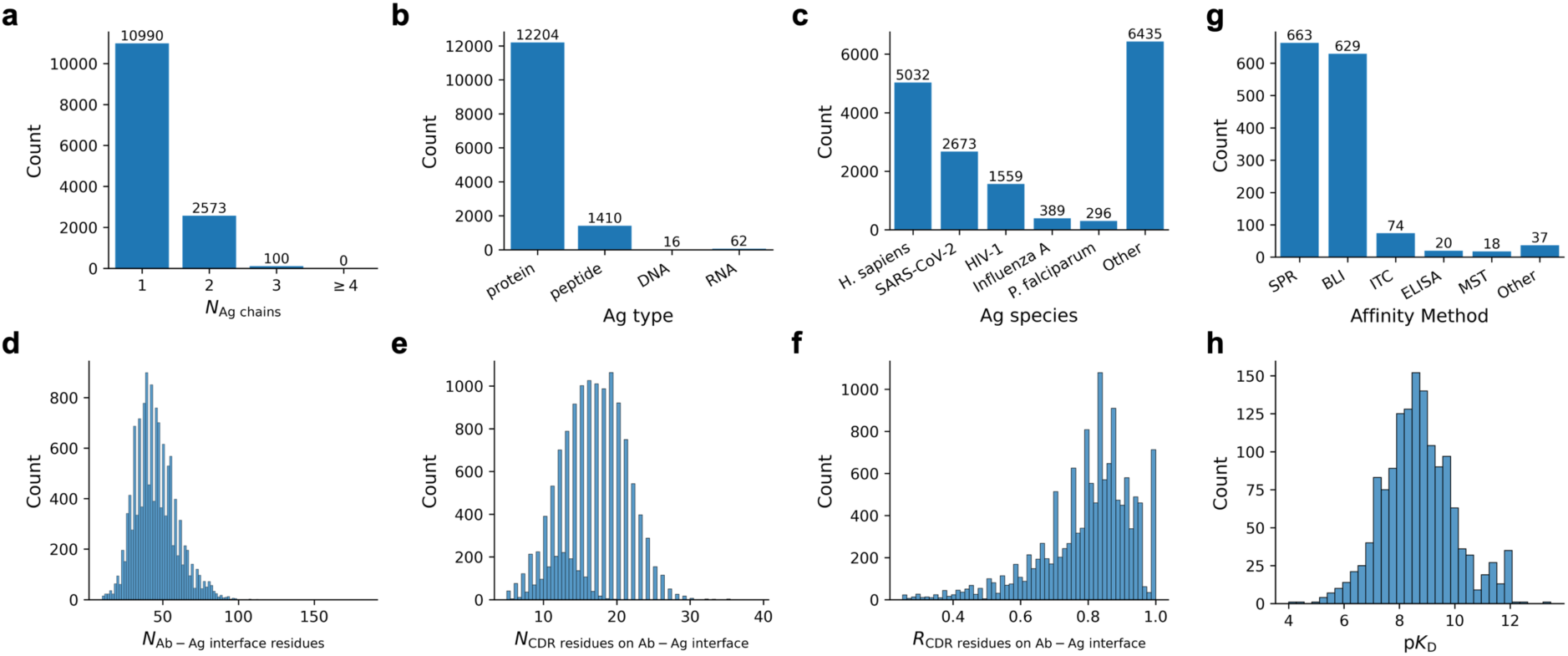
Statistics of antibody-antigen interactions (AAIs) in SAAINT-DB. **a**, Histogram of AAIs with varying numbers of antigen chains. **b**, Histogram of AAIs for different antigen types. **c**, Top sources of antigens. **d**, Histogram of the number of antibody-antigen interface residues. **e**, Histogram of the number of CDR residues on antibody-antigen interfaces. **f**, Histogram of the ratio of interfacial CDR residues to the total number of interface residues. **g**, Top methods for antibody-antigen binding affinity measurement. **h**, Histogram of p*K*_D_ values.

AAIs exhibit substantial diversity, with the number of residues at the Ab-Ag interfaces (*N_ab_ag_inf_res_*) ranging from 10 to 175, despite most falling between 30 and 60 (Fig. 3d). A key structural feature of AAIs is that Abs primarily engage Ags through their CDR residues. Accordingly, a significant portion of CDR residues is present at the Ab-Ag interfaces, with counts (i.e., *N_CDR_inf_res_* values) ranging from 5 to 40 (Fig. 3e), reflecting varied binding characteristics. The proportion of interfacial CDR residues to total interfacial Ab residues (*R_CDR_inf_res_*) spans from 25% to 100%, with most exceeding 70% (Fig. 3f), consistent with the knowledge that most interfacial Ab residues reside within the CDR regions.

In developing Ab therapeutics, a critical step is to optimize an Ab’s affinity for its target. Therefore, integrating Ab-Ag binding affinity data into SAAINT-DB is essential. For each PDB entry, we carefully reviewed its associated and relevant publications to extract the binding affinity data, if available. However, manually checking literature is laborious and very time-consuming. As of this writing, we have reviewed 2,866 out of the 9,373 PDB entries, 1,331 of which are associated with binding affinity data, accounting for 1,444 nonredundant data entries in SAAINT-DB-full.

The top five affinity measurement methods include surface plasmon resonance (SPR), biolayer interferometry (BLI), isothermal titration calorimetry (ITC), enzyme-linked immunosorbent assay (ELISA), and microscale thermophoresis (MST), with respective counts of 663, 629, 74, 20, and 18 entries, respectively, whereas all other methods contribute to 37 entries only (Fig. 3g).

Ab-Ag binding affinity, in terms of dissociation constant (*K*_D_), varies in a wide range from high micromolar to sub picomolar, with most in the nanomolar range, with p*K*_D_ values ranging from 4 to 14 with a median of ∼9 (Fig. 3h).

### Ab type classification

A major advancement of SAAINT-DB over existing structural Ab databases is its detailed classification of Ab types. Assessing classification accuracy is therefore crucial. Fig. 4 showcases structural examples of the eight most abundant Ab types in SAAINT-DB, demonstrating a high degree of consistency between algorithmic classification and structural visualization.

**Fig. 4.**
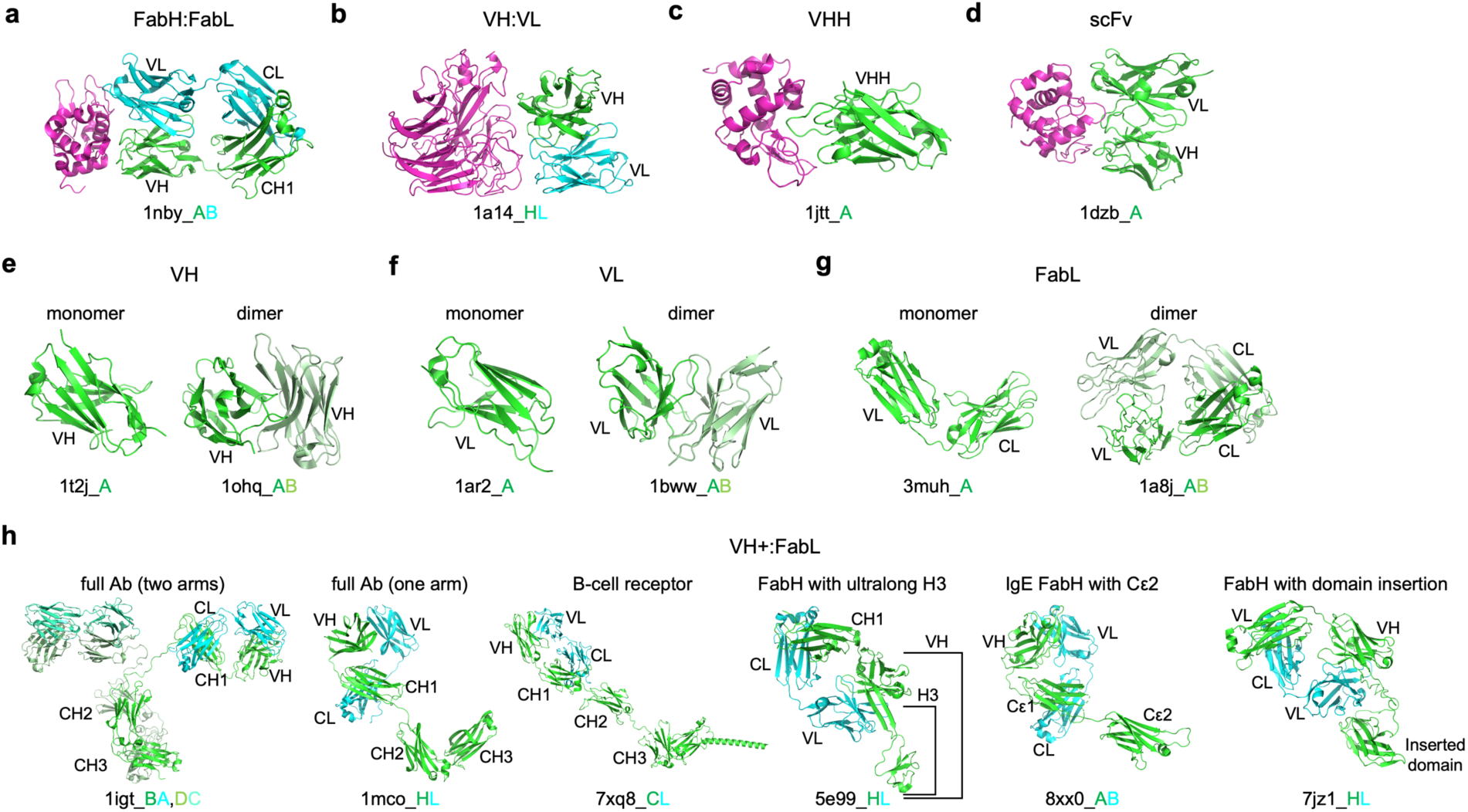
Example structures for the most abundant antibody types in SAAINT-DB. Antibody heavy chains, light chains, and antigen chains are shown in green, cyan, and magenta, respectively. Antibody domains such as VH, VL, CH1-3, and CL are labeled for clarification.

For instance, Fab 1nby consists of an HC and LC with VH-CH1 and VL-CL domains, respectively (Fig. 4a). Fv 1a14 features a VH:VL pair (Fig. 4b), while VHH 1jtt consists of a single VH domain from *Camelus dromedarius* (Fig. 4c). The scFv 1dzb represents a fused VH-VL construct (Fig. 4d).

Interestingly, certain types, such as VH, VL, and FabL, can exist either as monomers or homodimers (Fig. 4e,f,g). Notably, FabL dimers, also known as Bence-Jones proteins, consist of Ab LCs and are produced by abnormal plasma cells. These proteins are clinically significant as they are often associated with conditions like multiple myeloma and other plasma cell dyscrasias, indicating monoclonal gammopathy. However, to prioritize HC-LC pairings and discourage LC-LC and HC-HC pairings, SAAINT-parser splits these homodimeric pairings into monomers.

The VH+:FabL category is particularly complex, encompassing full-length Abs, B-cell receptors, Fabs with ultralong CDR-H3, IgEs with Cε2, and Fabs with domain insertions in the HC (Fig. 4h).

SAAINT-DB also accurately assigns Ab types even when Ab structures have numerous missing residues (Extended Data Fig. 1). For example, in VH-6pw6, only two CDR fragments are present in the HC, while scFv-6kn9 shows a significant number of missing residues. In VH-:VL-7ujd, both chains retain only three CDR loops. VHH-8jys has only two CDR fragments. In VH-:VL 7dk7, the VL is intact, but the VH contains just one CDR fragment. Similarly, in VH:VL-8vzo, the VH is complete, but the VL is notably incomplete.

Additionally, SAAINT-DB consistently and reliably categorizes Ab types in other cases as well (Extended Data Fig. 2).

### The SAAINT-DB representative dataset

Notably, many PDB entries contain multiple structural copies of the same Ab, making SAAINT-DB-full highly redundant. To address this, we developed a method to select the most representative Ab or AAI from the duplicated copies within each PDB entry by taking both Ab and Ag information into account (see Methods).

For example, in structure 3f12, where no Ag is present, the DC pair was chosen as the representative FabH:FabL because chains D and C have longer PDB-seq lengths and more interface residues than chains B and A (Fig. 5a). In structure 5yax, the B:C pair was selected as it includes the Ag peptide (chain C) (Fig. 5b). In structure 5zxv, both DC:B and HL:A pairs contain Ab and Ag; however, DC:B was chosen as the representative entry due to its larger Ab-Ag interface and a greater number of CDR residues at the interface (Fig. 5c).

**Fig. 5.**
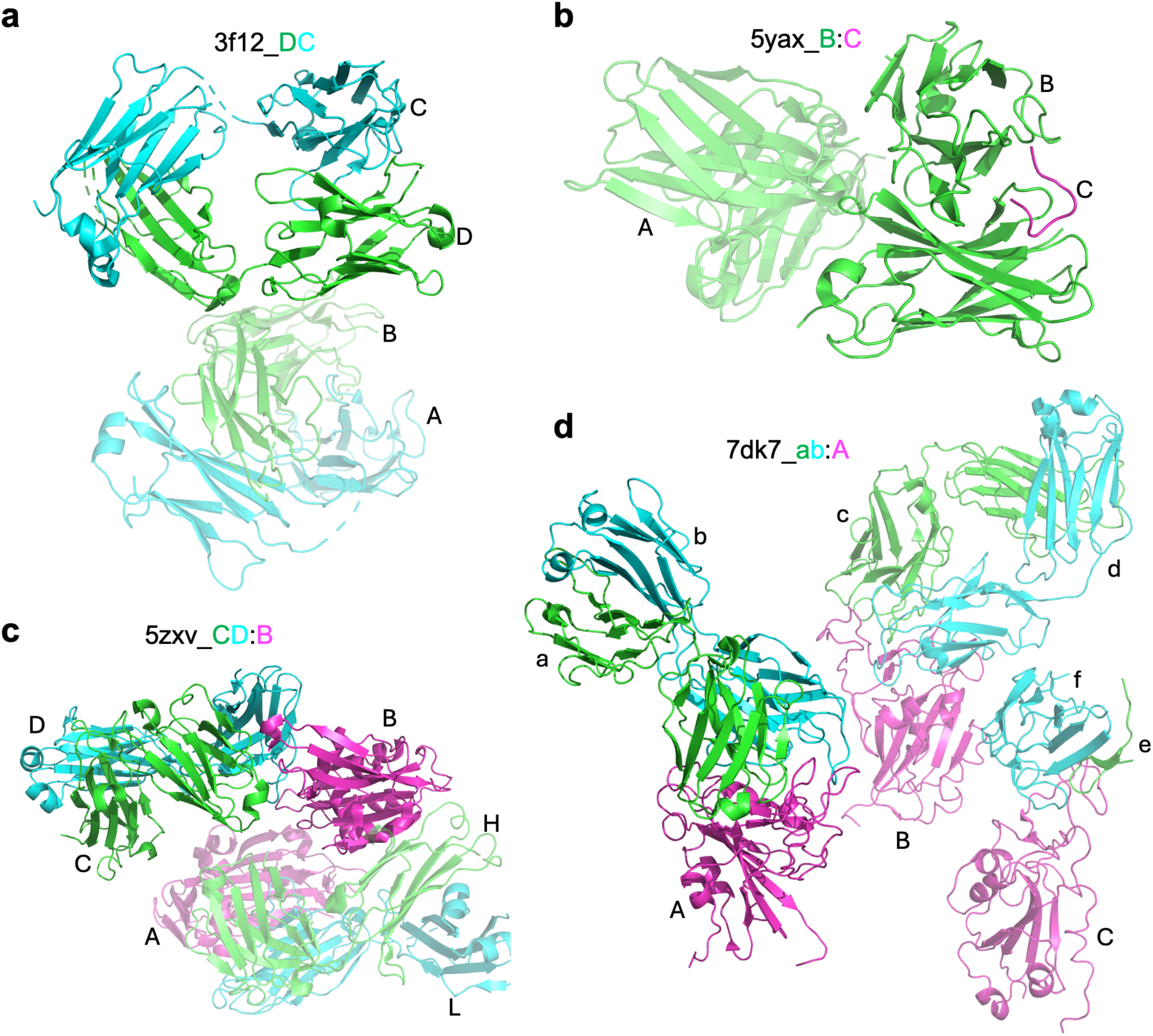
Example structures for representative antibody (Ab) and antibody-antigen interaction (AAI) entries in SAAINT-DB. Ab heavy chains, light chains, and antigen chains are shown in green, cyan, and magenta, respectively. In each case, the representative Ab or AAI is highlighted, while the non-representative Ab(s) and AAI(s) are shown with a cartoon transparency of 0.5.

The resulting dataset, SAAINT-DB-rep, consists of 10,199 entries, indicating approximately 43.4% (7,832/18,031) redundancy of SAAINT-DB-full at the PDB level. We conducted a similar statistical analysis of SAAINT-DB-rep to examine its distributions of Ab and AAI features (Extended Data Figs. 3, 4). Overall, these distributions closely resemble those of SAAINT-DB-full. FabH:FabL, VH:VL, VHH, scFv, VH, VL, FabL, and VH+:FabL remain the eight most abundant Ab types in SAAINT-DB-rep with 5,340, 1,708, 1,705, 851, 240, 97, 63, and 57 entries, respectively.

Filtering for representative Abs and AAIs slightly alters the number of Ab types. SAAINT-DB-full has defined 30 Ab types, while SAAINT-DB-rep has 29. The only Ab type that has been filtered out, VH-:VL (Extended Data Fig. 1, 7dk7 chains e,f), represents a low-quality, incomplete copy of the representative FabH:FabL Ab (Fig. 5d, 7dk7 chains a,b).

### Comparison with existing databases

The IMGT/3Dstructure-DB, last updated on May 23, 2024, contains 8,890 data entries and 7,243 PDB entries. AbDb, with its most recent update on July 26, 2019, includes 5,976 complete Ab Fvs and unpaired HCs or LCs derived from 3,348 PDB entries. The BEID and AgAbDb databases are currently inaccessible.

SAbDab, updated on January 17, 2025, has 18,265 data entries (representing Fv regions) from 9,200 PDB entries. Of these, 7,362 PDB entries contain at least one paired VH/VL, and 7,471 include Ags. Additionally, SAbDab provides 739 manually curated Ab-Ag affinity data points (Fig. 6a).

**Fig. 6.**
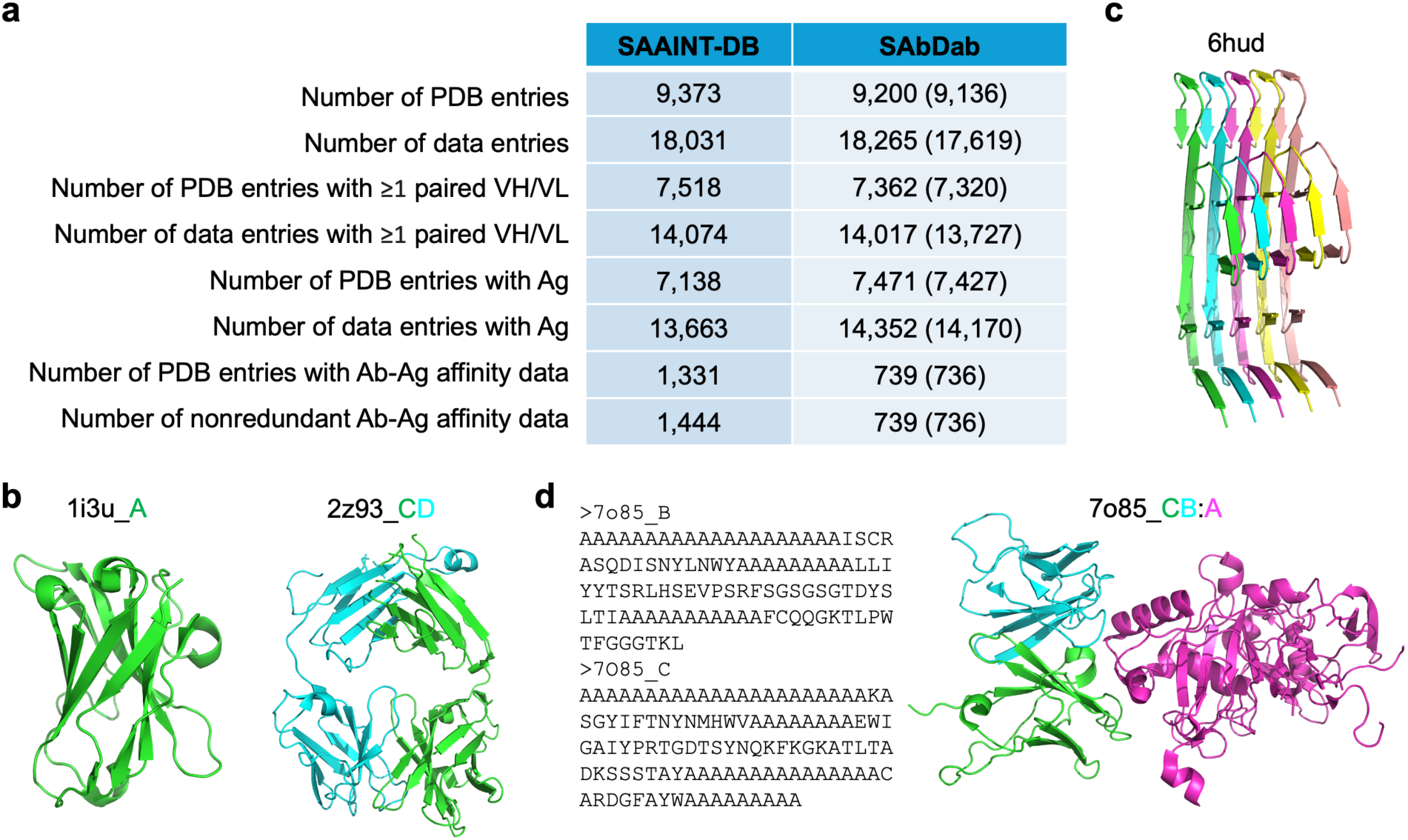
Comparison between SAAINT-DB and SAbDab. **a**, Statisical analyses of SAAINT-DB and SAbDab. **b**, Two examples of typical Ab structures recorded in SAAINT-DB but not in SAbDab. **c**, An example of amyloid fibrils from antibody light chain amyloidosis recorded in SAbDab but excluded from SAAINT-DB. **d**, An example of SAAINT-parser failure due to incorrect FASTA sequences.

In comparison, SAAINT-DB-full, last updated on January 29, 2025, comprises 18,031 data entries from 9,373 PDB entries. It includes 7,518 structures with at least one paired VH/VL, and 7,138 with Ags (Fig. 6a). Of the 2,866 out of 9,373 PDB entries reviewed, 1,444 nonredundant Ab-Ag binding affinity data points were collected, associated with 1,331 PDB structures.

To understand why SAAINT-DB contains more PDB entries but fewer Ab data entries than SAbDab, we investigated their differences in detail. Our analysis revealed that SAAINT-DB includes 239 PDB entries absent from SAbDab (Supplementary Table 1), such as the VHH A52 from *Lama glama* (PDB ID: 1i3u) and the anti-ciguatoxin Fab 10C9 (PDB ID: 2z93) (Fig. 6b). The reason for SAbDab’s exclusion of these typical Ab structures remains unclear. Conversely, SAbDab contains 66 PDB entries not present in SAAINT-DB, including 10 entries related to amyloid fibrils from Ig LC amyloidosis that lack typical Ab domains (Supplementary Table 2) (Fig. 6c, PDB ID: 6hud). One entry (PDB ID: 7o85) lacks correctly assigned Ab FASTA-seqs despite being annotated as containing a Fab by the deposition authors (Fig. 6d). Another entry (PDB ID: 2h3n) represents the structure of a surrogate LC homodimer; its LC FASTA-seqs were unprocessable by AbRSA due to a large number of missing residues at the C-terminus, although its companion entry (PDB ID: 2h32) was processed successfully (Extended Data Fig. 5). The remaining 54 entries are either removed or have been replaced in the PDB (Supplementary Table 3), leading to their exclusion from SAAINT-DB.

The two databases also differ in their treatment of shared entries. SAbDab generates an Ab entry for each model within an assembly (Supplementary Table 4), resulting in a large number of duplicated entries for a single PDB entry, whereas SAAINT-DB creates only one entry for each assembly. For example, SAbDab records 77 Ab entries for PDB entry 2kh2, while SAAINT-DB has only one. Additionally, SAbDab failed to pair Abs in 60 PDB entries which were faithfully paired in SAAINT-DB (Supplementary Table 5).

After excluding the obsolete PDB entries and duplicated NMR models, a refined SAbDab would contain 17,619 data entries derived from 9,136 PDB structures (Fig. 6a). Overall, SAAINT-DB outperformed SAbDab on most metrics except for the number of PDB and data entries with Ag (Fig. 6a), as SAAINT-DB currently excludes carbohydrate and hapten Ags.

## Discussion

While several specialized structural Ab databases exist, they still have notable limitations. This study aims to develop a comprehensive and accurate database that can complement or potentially improve over existing resources. To achieve this, we developed SAAINT-parser, the first open-source workflow for fast and accurate parsing of PDB entries to extract structural Abs and AAIs. A detailed comparison with SAbDab and other databases highlights the advantages of the resulting SAAINT-DB in both accuracy and completeness.

Accurate pairing of HCs and LCs is a crucial step in SAbDab, AbDb, and SAAINT-DB. This process relies not only on the precise identification of Ab chains but also on correctly matching corresponding HCs and LCs. We used AbRSA to identify Ab chains and CDR regions, leveraging its high accuracy and speed^20^. Existing databases rely on different strategies for HC-LC pairing. SAbDab enforces a constraint requiring the conserved cysteine at Chothia position 92 on an HC to be within 22 Å of the conserved cysteine at position 88 on an LC^15^. While effective in most cases, this criterion is overly restrictive for certain structures and fails when one or both of these cysteines are missing, which may partly explain why SAbDab sometimes identifies Ab chains that should be paired as unpaired (Supplementary Table 5). AbDb, on the other hand, maximizes the number of atomic contacts within 4 Å to determine pairings^19^. While this approach works well for single-domain structures (e.g., VHs and VLs) in building the AbDb database, it can lead to incorrect pairings in complex cases such as domain-swapped Fabs.

To overcome these challenges, SAAINT-parser implements a robust HC-LC pairing procedure. First, we defined thresholds for *N_HL_inf_res_* to filter out false pairings while improving computational efficiency. Second, we prioritized high-confidence pairings using a composite scoring function that combines *N_HL_inf_res_* (similar to AbDb’s contact-based approach) with the mean index of interface residues. Finally, we employed a hybrid approach combining greedy search and iterative heuristic search to maximize the number of valid HL pairs across the entire structure. This method ensures accurate pairings even in very complicated cases. For example, in PDB entry 8d01, which contains two HCs (A and H) and two LCs (B and L), the number of interface residues for pairings AB, AL, HB, and HL are 44, 50, 52, and 48, respectively. AbDb’s rule predicts AL and HB as the correct pairings, whereas the actual pairings are AB and HL (Extended Data Fig. 6a). A more complex example is PDB entry 2oqj, which consists of four HCs (B, E, H, and K), four LCs (A, D, G, and J), and four Ag chains (C, F, I, and L). The number of interface residues for HC-LC pairings BA, EA, BD, ED, HG, KG, HJ, and KJ are 51, 60, 56, 50, 48, 57, 55, and 50, respectively. Under AbDb’s rule, the predicted pairings would be EA, BD, KG, and HJ, while the correct pairings are BA, ED, HG, and KJ (Extended Data Fig. 6b).

Regular updates are essential for any database. SAbDab is well updated with the latest update on January 17, 2025. In contrast, the IMGT/3Dstructure-DB and AbDb databases are updated less frequently, with their most recent updates on May 23, 2024 and July 26, 2019, respectively. Our tests showed that scanning the entire PDB database with 300 CPUs (3.0 GHz Intel Xeon Gold 6154) took only a few hours to build the initial SAAINT-DB datasets and even less time for updates (see Methods). Therefore, we conclude that SAAINT-DB could be consistently kept up to date with the PDB.

Despite its advantages, SAAINT-parser and SAAINT-DB have certain limitations. First, SAAINT-parser relies on AbRSA for Ab chain type identification, making its accuracy dependent on AbRSA’s precision. While AbRSA is highly accurate, it can occasionally fail. For example, in PDB entry 8d53, AbRSA failed to identify the HC of 35022scFv, likely due to a missing nine-residue fragment in its FASTA-seq. However, AbRSA and SAAINT-parser successfully identified other Ab chains, including Fab PGT124. Notably, this entry is completely absent from SAbDab. Second, classifying engineered or unusually long Ab chains remains somewhat ambiguous. For instance, VH+ indicates the presence of a VH domain but does not specify the exact number of VH domains or provide structural details of non-VH regions. Similarly, VH+:FabL encompasses diverse Ab structural configurations (Fig. 4h). Third, SAAINT-parser and SAAINT-DB currently support only protein, peptide, RNA, and DNA Ags, limiting their applicability to other Ag types such as carbohydrates and haptens (nonpolymeric ligands). Addressing these limitations is a priority for future development.

## Methods

### Data source

The mmCIF files of the whole PDB repository were downloaded to a local high-performance computing cluster following the wwPDB guidelines at https://www.wwpdb.org/ftp/pdb-ftp-sites. Each entry’s corresponding FASTA file and PDB information were retrieved from https://www.rcsb.org/fasta/entry/pdbid and https://www.rcsb.org/structure/pdbid, respectively, where pdbid represents a valid PDB ID.

### FASTA and mmCIF data structure

For each PDB entry, the FASTA file contains sequences for all macromolecular entities recorded in the mmCIF file. Each entity consists of one or more chains with ID(s) labeled in label_asym_id (assigned by the PDB) and often auth_asym_id (designated by the deposition authors). All chains within an entity share the same sequence (FASTA-seq). The entity also includes a molecule name and the species from which it originates.

The mmCIF file provides atomic coordinates for both the entities recorded in FASTA and other unrecorded molecules, such as ions, cofactors, water molecules, and nonpolymeric ligands. In this work, we focused on the structures of Abs and their interactions with proteins, peptides, and nucleic acids while ignoring other molecules. Although the mmCIF file encodes all the information found in the FASTA file, both formats were used because FASTA, being simpler, enables more efficient data processing.

### Identifying and labeling Ab chains

Each FASTA-seq was analyzed using AbRSA^20^ to determine whether it contains an Ab chain. Based on this analysis, each FASTA-seq was categorized into one of the following types: HC, LC, HLC, or non-Ab. If all chains were labeled as non-Ab, the entry was deemed to lack an Ab chain and was discarded.

For entries containing at least one Ab chain, each protein or peptide structure was extracted and saved as a separate PDB file using Biopython^21^. The corresponding primary sequence (PDB-seq), composed of residues with atomic coordinates, was then derived. To ensure accurate sequence mapping, the PDB-seq was aligned to the FASTA-seq using Biopython’s Align module with customized parameters (Supplementary Table 6). Based on this alignment, a gap-filled PDB-seq (Filled-seq) was generated, spanning the minimum to maximum aligned positions. The Filled-seq was then subjected to a second AbRSA analysis to examine the presence of an Ab chain within the structure.

The rationale for using Filled-seq in AbRSA type labeling is that it more closely reflects the actual structure compared to FASTA-seq while incorporating missing residues to ensure sequence continuity compared to PDB-seq. This approach is particularly useful in cases where the FASTA-seq contains an Ab sequence, but the corresponding atomic coordinates are absent in the structure. Such structures were excluded from SAAINT-DB due to their incompleteness in Ab structures.

If a Filled-seq was identified as Heavy or Light, it was further analyzed using Abalign^27^ to determine its V gene subgroup. However, sequences labeled as HeavyLight were excluded from subgroup analysis, as their hybrid nature—either as a scFv from the same species or a chimera of VH and VL domains from different species— could lead to inaccurate predictions.

Some structures, despite containing Ab chains, lack typical Ab structural domains. For instance, AL amyloid fibrils result from the misfolding of Ab LCs (e.g., PDB ID: 7nsl^28^). To exclude such cases, TM-align^29^ was used to compare these structures against high-quality reference domains—VH (PDB ID: 6cnw^30^, chain A, X-ray resolution: 0.92 Å) or VL (PDB ID: 4unu^31^, chain A, X-ray resolution: 0.95 Å)—based on their AbRSA classification as HC or LC. Ab chains with a TM-score^32^ (normalized to the reference) below 0.4 were discarded. Notably, all LCs in 7nsl exhibited TM-scores below 0.2 when compared to 4unu, confirming their lack of Ab domains.

### Pairing HCs, LCs, and HLCs

Once AbRSA types were assigned, Ab chains were paired based on their level of interaction. However, structural limitations such as extensive missing residues/atoms or models containing only Cα atoms could hinder accurate interaction calculations. To address this, we used Pulchra^22^ to reconstruct full-atom protein models from the Cα traces or incomplete structures. During reconstruction, nonstandard amino acids and residues lacking Cα atoms were excluded, as Pulchra could not process them correctly. The repaired Ab chain structures were then analyzed using AbRSA_PDB^20^ to identify CDR residues. Finally, all the repaired chains were merged into a single complex structure and subjected to side-chain repacking using FASPR^23^ to minimize clashes.

Ab chains were then paired both within and between entities using UniDesign^24,25^ to compute the number of interface residues between potential pairing chains (*N_HL_inf_res_*). A residue was considered part of the interface if at least one of its nonhydrogen atoms was within 5 Å of a nonhydrogen atom in the other chain^33,34^. Typically, a VH-containing chain was paired with a VL-containing chain. Pairing HLCs was more complex, as these chains contain both VH and VL and could pair with HCs, LCs, or other HLCs.

To ensure accurate pairings, a variety of minimum *N_HL_inf_res_* cutoffs were applied, based on chain lengths (Supplementary Table 7). For example, two scFv chains (both are HLCs) were considered a valid pair if their *N_HL_inf_res_* was at least 80, whereas a VH-VL pairing (HC and LC) required only ≥20 interface residues. This filtering step effectively excluded chains that were in proximity due to crystal packing but did not form a true functional interface.

Under these criteria, one chain may still structurally pair with multiple partner chains. However, each chain should ideally pair with only one other chain or remain unpaired. To resolve cases where an HC pairs with multiple LCs or vice versa, we employed a combination of greedy search and iterative heuristic search, guided by a carefully designed score function, to maximize the number of valid HL pairs within the entire structure.

In the greedy search procedure, the initial HC-LC pairings were determined using the following steps: (1) Each unpaired Ab chain was temporarily paired with a dummy chain to form a placeholder HL pair. (2) All HL pairs were scored (see the following subsection) and ranked from the highest to lowest based on their pairing scores. (3) A greedy HC-LC pairing solution was constructed by iteratively selecting the highest-scoring HL pairs, ensuring that each chain was only used once. This process continued until all real Ab chains were covered. The total greedy score was then computed by summing the pairing scores of the selected HL pairs. (4) Unused HL pairs were pooled for further evaluation.

The initial pairing solution by the greedy search may not be optimal. To further refine the pairings, an iterative heuristic search was performed as follows: (1) Starting from the greedy pairing solution, the highest-scoring HL pairs were progressively replaced with unused HL pairs to construct a new HC-LC pairing solution while ensuring that each chain was only used once. (2) A new total pairing score was calculated for the updated solution. (3) The substituted HL pairs were returned to the unused pool. (4) This process was repeated until replacing pairs no longer resulted in a higher total score.

This combined strategy ensured a more accurate and stable HC-LC pairing assignment than the greedy search alone, resolving ambiguities and optimizing the structural pairing of Ab chains.

### Score function for HC-LC pairing

Each HL pair was scored using the following scoring function:

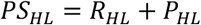

where *R_HL_* and *P_HL_* represent the reward and penalty terms for HC-LC pairing, respectively. These terms are defined as follows:

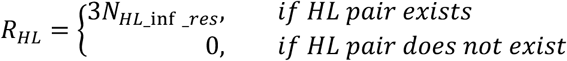

where *N_HL_inf_res_* denotes the number of interface residues between HC and LC.

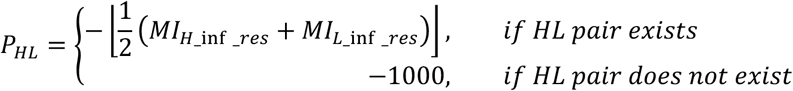

where *MI_H_inf_res_* and *MI_L_inf_res_* represent the mean positions of interface residue on HC and LC, respectively.

### Calculating Ab chain mean radius

The mean radius of an Ab chain was calculated using the following equation:

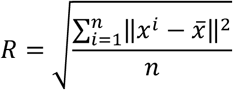

where *n* is the number of heavy atoms in the chain, *x^i^* represents the coordinates of atom *i*, and *x̄* denotes the averaged coordinates of all heavy atoms.

### Determining Ab types

The Ab type classification was determined based on multiple factors, including the AbRSA type, molecular name, species name, mean radius, and the lengths of PDB-seq, Filled-seq, and FASTA-seq. Briefly, an HC was classified into one of the following types: FabH, VH, VH+ (an HC longer than a typical VH), VH-(an HC shorter than a typical VH), VHH, VHH+, and VHH-. In contrast, an LC was categorized as FabL, VL, VL+, or VL-. LCs do not include VHH, VHH+, or VHH-types, as these are exclusive to HC-only Abs. An HLC was classified as scFv, scFv+, or scFv-if its molecular name contained keywords such as *L*single,*L L*scFv,*L* or *L*Fv*L* and its mean radius was ≤20 Å. Otherwise, it was categorized as VHVL (a VH-VL fusion of typical scFv size but not explicitly identified as scFv), VHVL+, or VHVL-.

### Pairing Ab and Ag chains

Each Ab was systematically paired with each Ag chain using the same approach as for HC-LC pairing. Specifically, a temporary Ab-Ag complex was constructed, and UniDesign^24,25^ was used to identify its interface residues. An Ag chain was classified as a true Ag if it met all three of the following conditions: (1) the number of Ab-Ag interface residues (*N_ab_ag_inf_res_*) was ≥10; (2) the number of CDR residues at the interface (*N*_-*CDR*_*__inf_res_*) was ≥5; and (3) the ratio of the number of interface CDR residues to the number of interface Ab residues (*R_CDR_inf_res_*) was ≥0.25. All Ag chains that met these criteria were grouped together, meaning that each Ab could interact with one or multiple Ag chains.

### Selecting representative Abs and AAIs

For PDB entries containing multiple copies of the same Ab or AAI, the most representative structure was selected using the following scoring function:

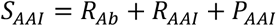

where *R_AB_* and *R_AAI_* are reward terms for Ab and AAI, respectively. *P_AAI_* is a penalty term applied when an AAI involves multiple Ag chains. These terms are defined as follows:

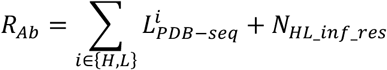

where *L_PDB-seq_* represents the PDB-seq length of HC or LC, and *N_HL_inf_res_* is the number of interface residues between HC and LC.

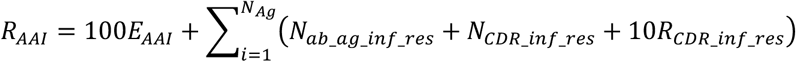

where *E_AAI_* is assigned a value of 1 if an AAI exists, otherwise 0. *N_Ag_* denotes the number of Ag chains. *N_ab_ag_inf_res_* is the number of Ab-Ag interface residues. *N_CDR_inf_res_* is the number of CDR residues at the interface. *R_CDR_inf_res_* is the ratio of CDR interface residues to total Ab interface residues.

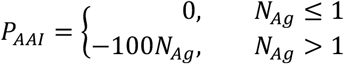

Copies of Abs or AAIs with the highest *S_AAI_* scores were selected as the most representative structures and recorded in SAAINT-DB-rep. As indicated, this scoring function prioritizes structures with higher structural integrity, more extensive and correctly oriented packing/binding, and a single Ag chain, while discouraging cases with multiple Ag chains.

### Ab-Ag binding affinity data

To ensure accurate integration of experimentally determined Ab-Ag binding affinity data into SAAINT-DB, each PDB entry was carefully cross-referenced with the PDB-associated publication and related references. Binding affinity data, presented as the dissociation constant (*K*_D_) in units of nM, was reported when available. Overall, various techniques were used to determine binding affinity, including surface plasmon resonance (SPR), biolayer interferometry (BLI), isothermal titration calorimetry (ITC), enzyme-linked immunosorbent assay (ELISA), kinetic exclusion assay (KinExA), solution equilibrium titration (SET), flow cytometry, and interferometry. When different methdos were used for an AAI, ITC data was prioritized for its accuracy. If multiple values were obtained using the same technique (e.g., SPR), the geometric mean was calculated. The temperature of the affinity measurement was recorded if explicitly stated, standardized to 298 K for room temperature, or marked as *L*N.A.*L* if unspecified.

### Recording Ab and AAI information

For each PDB entry identified as containing an AAI or Ab, essential information was extracted and recorded, encompassing a total of 42 attributes related to the Ab, Ag, and PDB (Supplementary Table 8). For unpaired HCs (e.g., scFv) without an LC partner, LC-related fields were assigned either *L*N.A.*L* or *L*0,*L* depending on the data type. Similarly, for unpaired LCs (e.g., VL), HC-related fields were marked as *L*N.A.*L* or *L*0.*L* If an Ab was not paired with an Ag, the Ag-related fields were also set to *L*N.A.*L* or *L*0.*L* All other unavailable fields were recorded as *L*N.A.*L*

### Building and updating SAAINT-DB

To construct SAAINT-DB for the first time, all mmCIF files from the PDB were downloaded to a local cluster using remote synchronization (rsync), while the associated FASTA files were retrieved via wget. The PDB entries were distributed across hundreds of CPUs, with each CPU processing <1,000 entries using SAAINT-parser. Based on our tests, the initial processing on 300 CPUs took approximately ten hours to complete. Once processed, the processed Ab and AAI data entries for individual PDB entries were merged to generate the SAAINT-DB full and representative datasets.

To update SAAINT-DB, rsync was used to detect removed, modified, and new added mmCIF files, and the corresponding FASTA files were updated accordingly. The Ab and AAI data entries were removed for deleted PDB entries, recomputed for modified ones, and newly computed for newly added entries. Only these updated PDB entries were assigned to a smaller number of CPUs, each running SAAINT-parser to process a few entries. Our tests showed that the update process typically took less than two hours, depending on the number of updated PDB entries. Finally, the processed Ab and AAI data entries were merged to update the SAAINT-DB-full and SAAINT-DB-rep.

Since Ab-Ag binding affinity data must be manually collected and curated from the literature, it lagged behind the SAAINT-DB datasets and was maintained separately.

## Data availability

The SAAINT-DB-full, SAAINT-DB-rep, and manually curated Ab-Ag binding affinity data are summarized in Excel sheets and are available at https://github.com/tommyhuangthu/SAAINT. Computed Ab and AAI structures are available upon request. The complete PDB repository can be downloaded from https://files.wwpdb.org/.

## Code availability

The SAAINT-parser source code is available at https://github.com/tommyhuangthu/SAAINT. It is built on Python 3.10 and Biopython 1.84. FASPR (v.20200309) can be accessed at https://github.com/tommyhuangthu/FASPR. UniDesign (v.1.2) is available at https://github.com/tommyhuangthu/UniDesign. AbRSA (v.2.20) can be downloaded from http://cao.labshare.cn/AbRSA/download.html, while AbRSA_PDB (v.2.00) is available through Dr. Yang Cao at Sichuan University. Abalign (v.2.00) can be downloaded from http://cao.labshare.cn/abalign/. TM-align (v.20220412) is available at https://zhanggroup.org/TM-align. Lastly, Pulchra (v.3.06) can be accessed at https://github.com/euplotes/pulchra.

## Acknowledgments

This work was supported by the Universitity of Michigan Medical School internal funds to the Center for Advanced Models for Translational Sciences and Therapeutics (CAMTraST). We thank the Advanced Research Computing (ARC) at the University of Michigan for providing the computational resources and services that supported this research.

## Author contributions

X.H. conceived and conducted the project. X.H., J.Z. S.C. X.X., Y.E.C. and J.X. conducted validation of the database. X.H. wrote the paper. X.H., Y.E.C. and J.X. edited the paper. All authors participated in discussions and agreed with the final paper.

---

**Extended Data Fig. 1.**
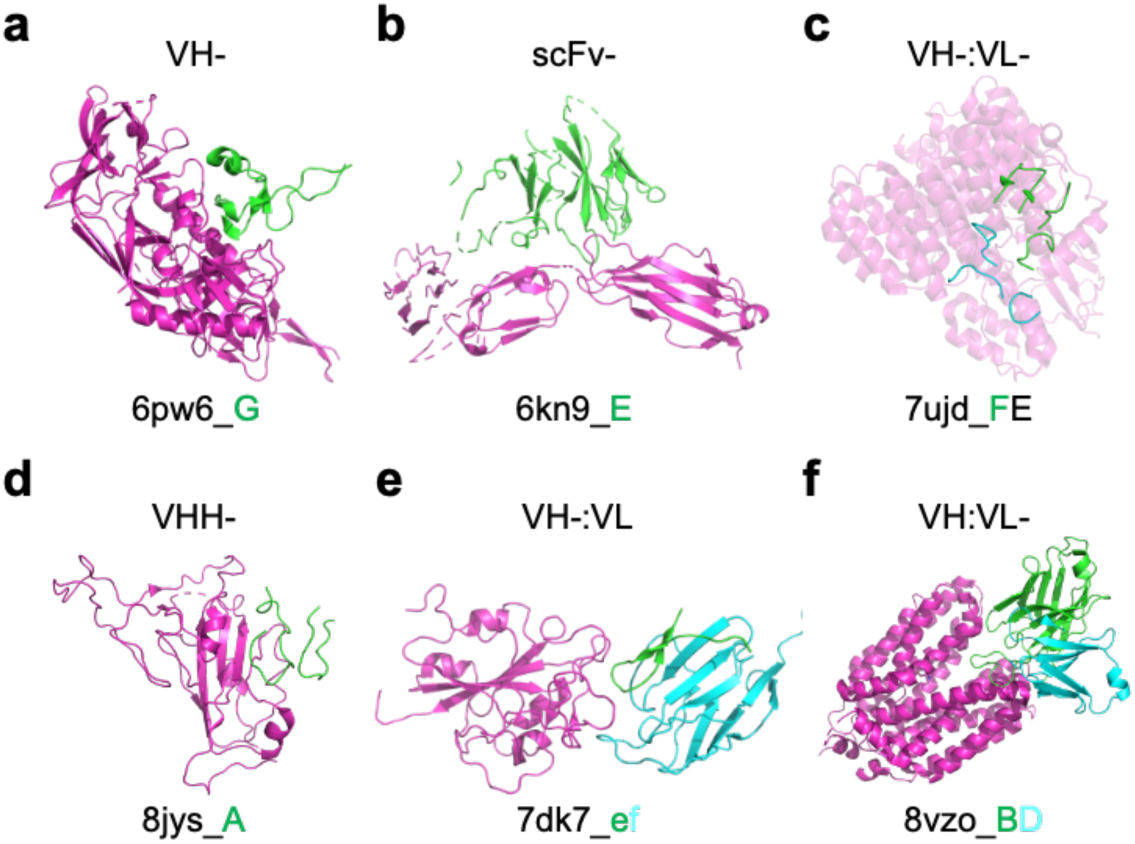
Example structures for antibody types with missing residues. Antibody heavy chains, light chains, and antigen chains are shown in green, cyan, and magenta, respectively.

**Extended Data Fig. 2.**
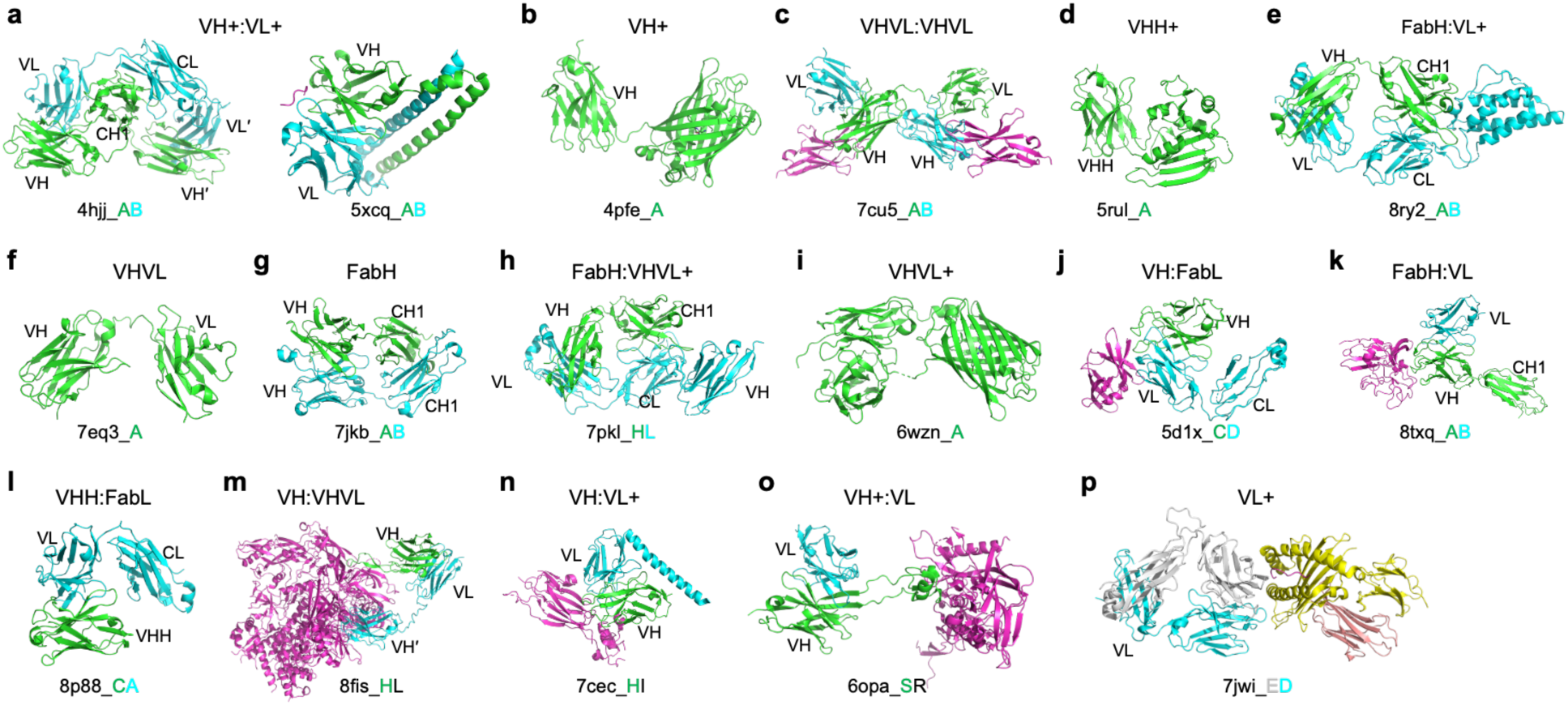
Example structures for other uncommon antibody types. Antibody heavy chains, light chains, and antigen chains are shown in green, cyan, and magenta, respectively, except for the VL+ type. Ab domains, such as VH, VL, and CH1 are labeled for clarification.

**Extended Data Fig. 3.**
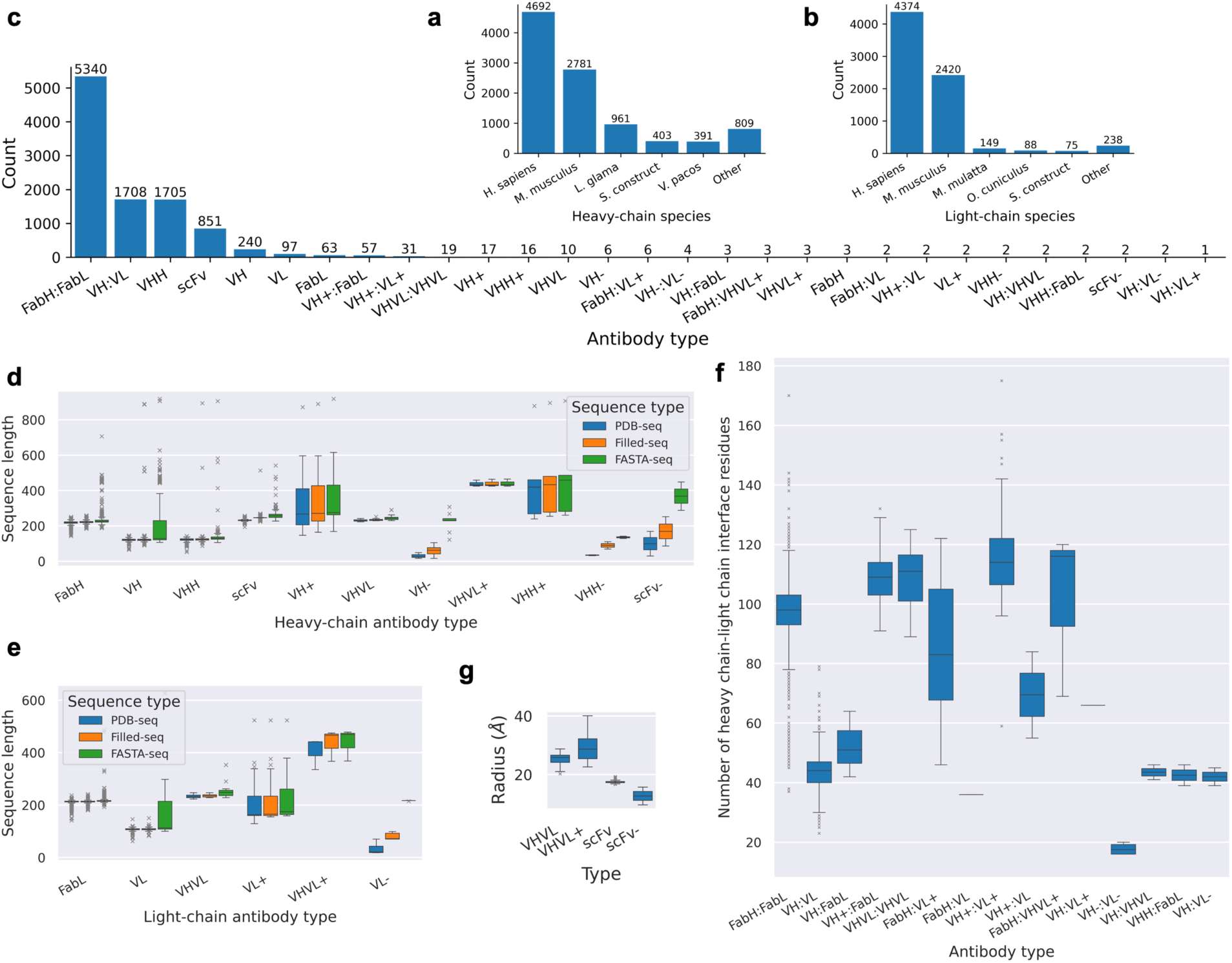
Statistics of antibodies in the SAAINT-DB representative dataset. **a**, Top sources of antibody heavy chains. **b**, Top sources of antibody light chains. **c**, Antibody type classification in SAAINT-DB-rep. **d**, Distrition of sequence lengths for heavy-chain antibody types. **e**, Distribution of sequence lengths for light-chain antibody types. **f**, Distribution of the number of heavy chain-light chain interface residues across distinct antibody types. **g**, Distribution of the mean radius for scFv and VHVL types.

**Extended Data Fig. 4.**
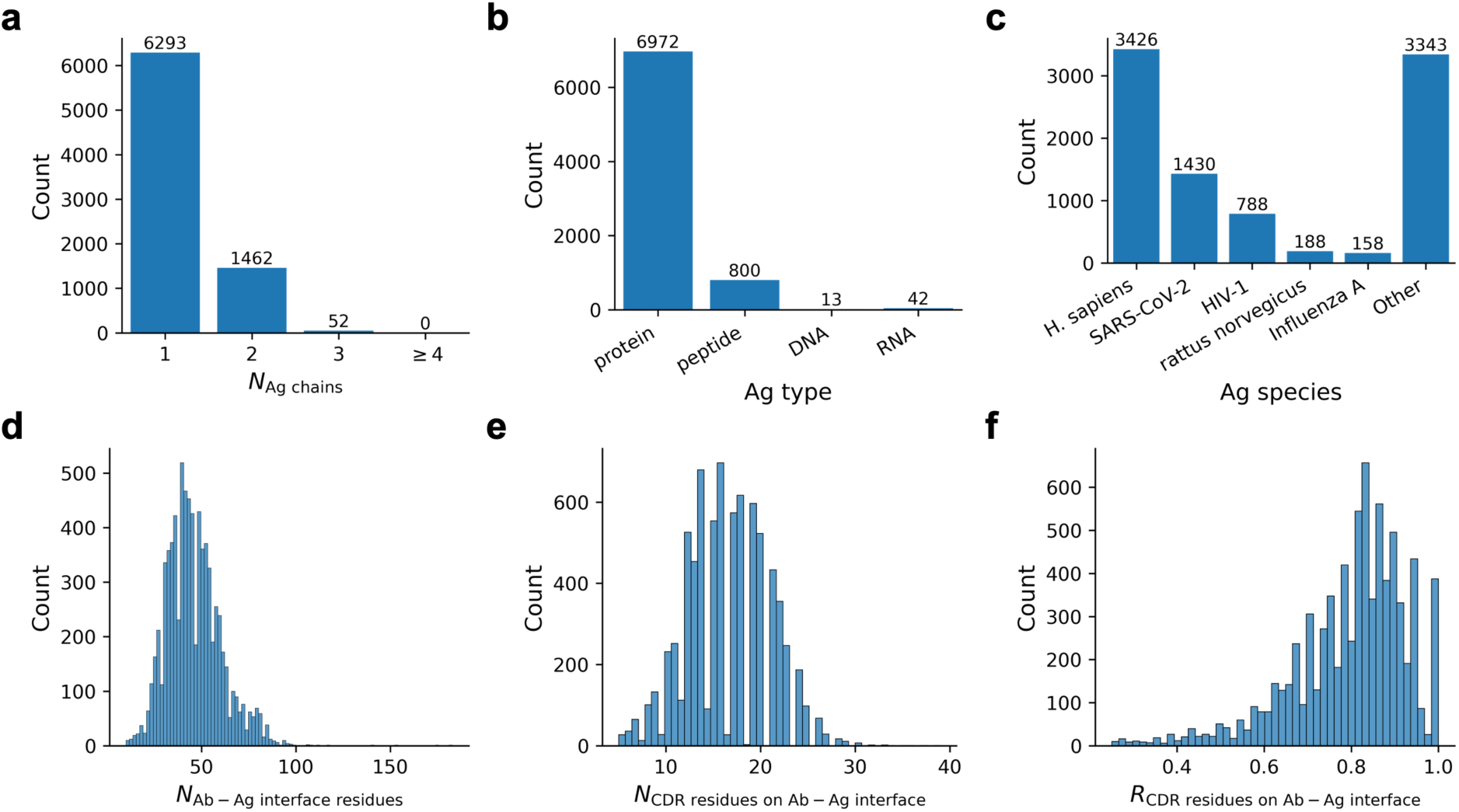
Statistics of antibody-antigen interactions (AAIs) in the SAAINT-DB representative dataset. **a**, Histogram of AAIs with varying numbers of antigen chains. **b**, Histogram of AAIs for different antigen types. **c**, Top sources of antigens. **d**, Histogram of the number of antibody-antigen interface residues. **e**, Histogram of the number of CDR residues on antibody-antigen interfaces. **f**, Histogram of the ratio of interfacial CDR residues to the total number of interface residues.

**Extended Data Fig. 5.**
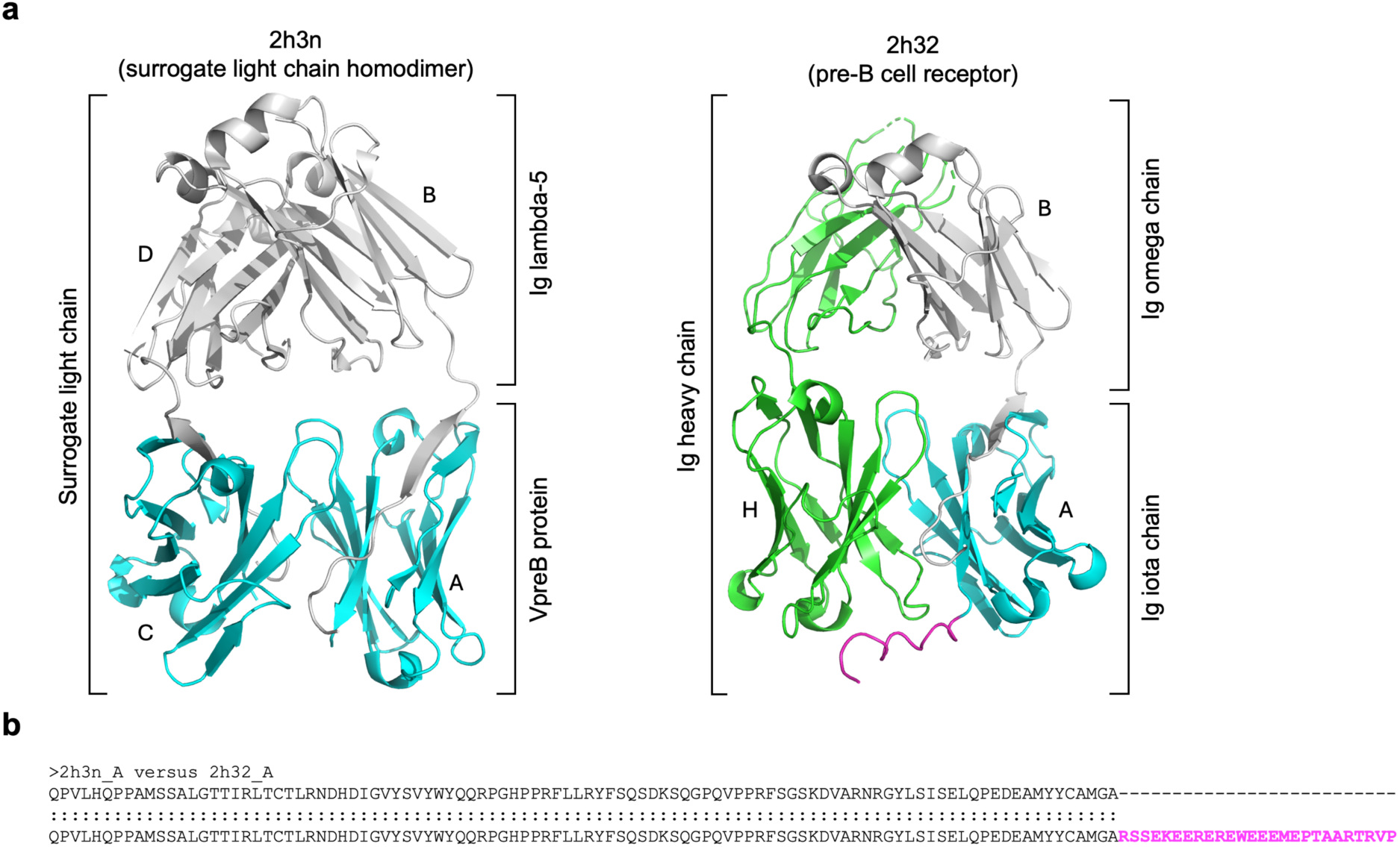
An example of SAAINT-parser failure. **a**, Structure comparison between 2h3n and 2h32. Heavy and light chains are shown in green and cyan, respectively, while other chains are shown in silver. **b**, Sequence alignment between 2h3n chain A and 2h32 chain A. The C-terminal residues absent in 2h3n chain A but present in 2h32 chain A are highlighted in magenta.

**Extended Data Fig. 6.**
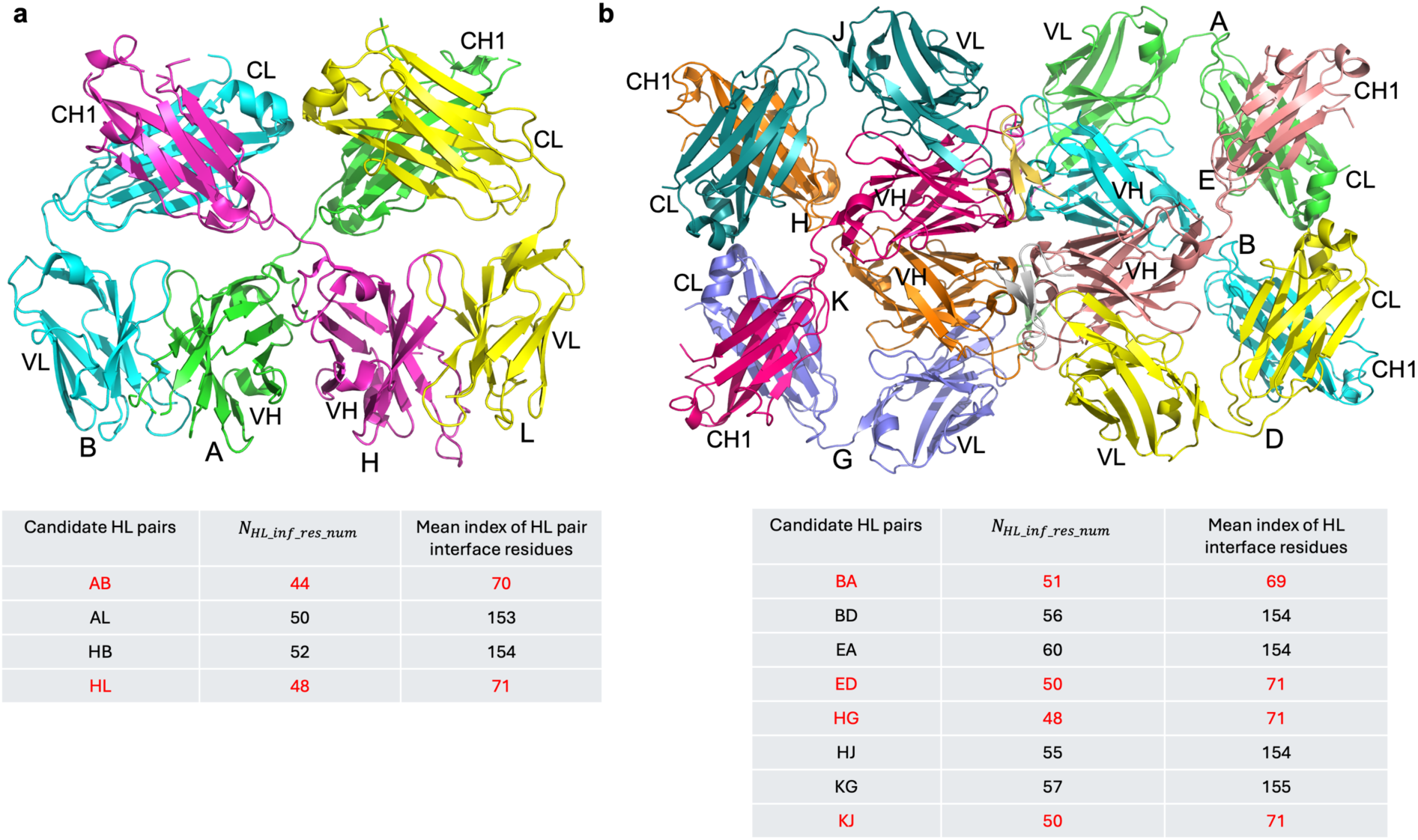
Examples of accurate Ab heavy and light chain pairings in SAAINT-parser. Correct heavy chain-light chain pairings are highlighted in red in the tables below.

## Supplementary information

**Supplementary Table 1.**
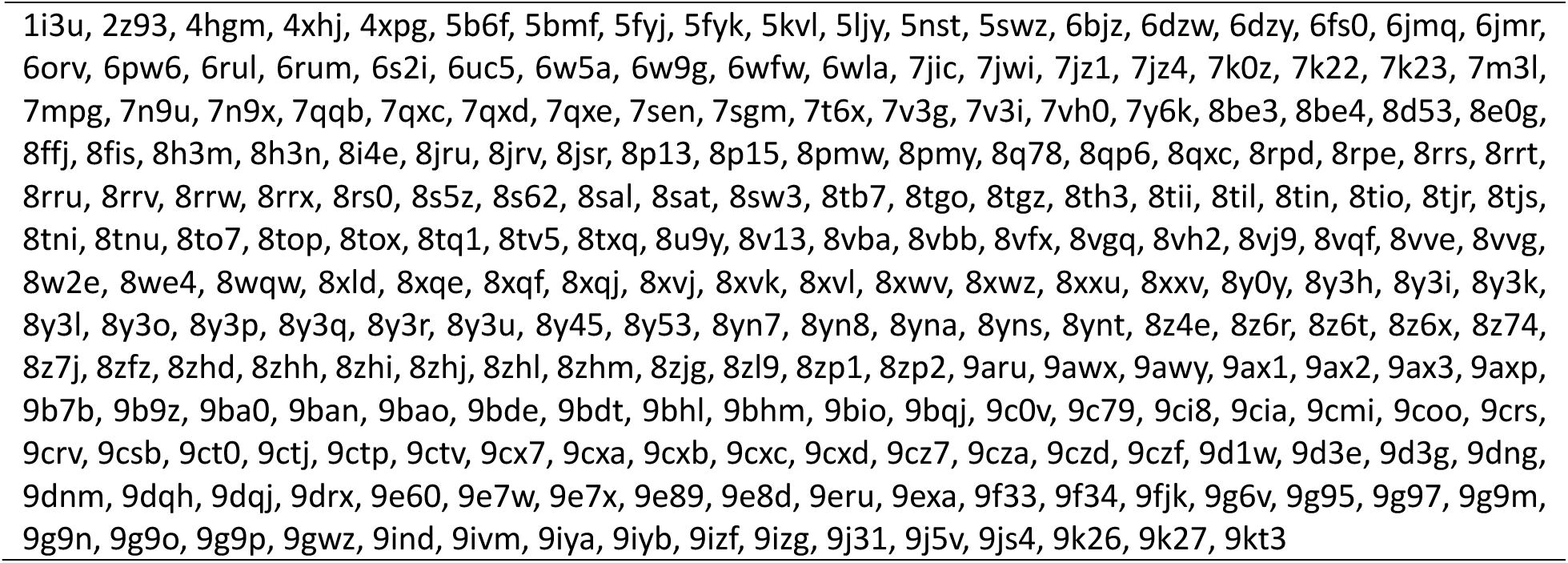
List of 239 PDB entries recorded in SAAINT-DB but not in SAbDab.

**Supplementary Table 2.**
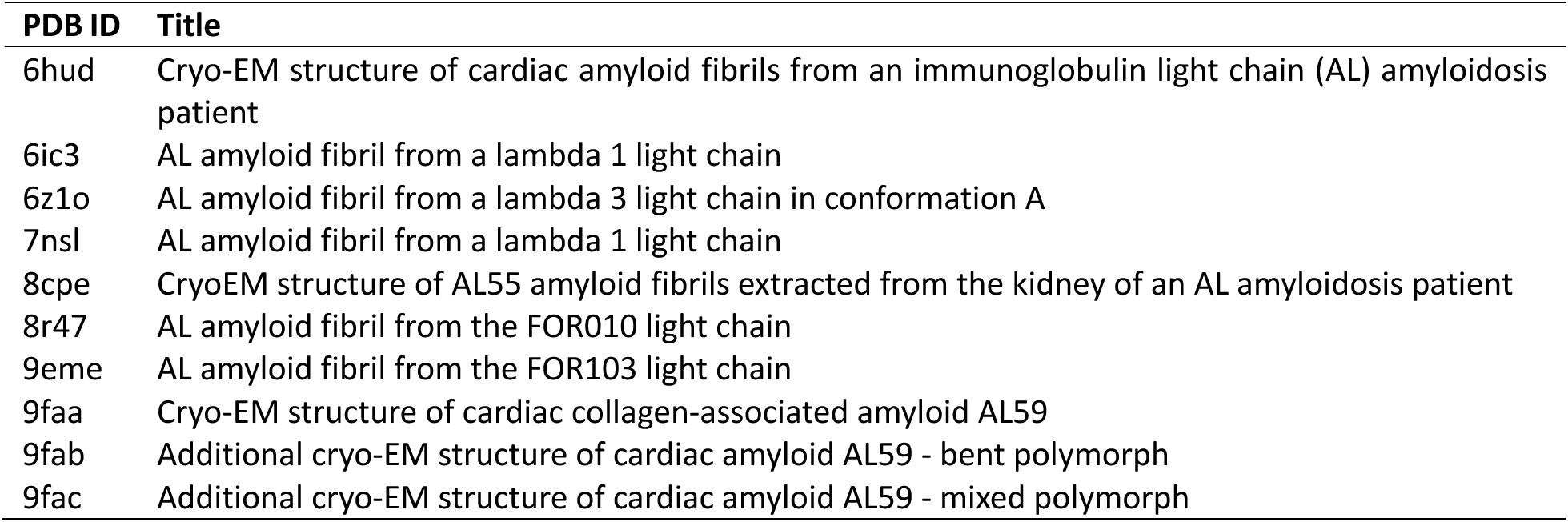
List of 10 PDB entries of antibody light chains forming amyloid fibrils.

| PDB ID | Title |
| --- | --- |
| 6hud | Cryo-EM structure of cardiac amyloid fibrils from an immunoglobulin light chain (AL) amyloidosis patient |
| 6ic3 | AL amyloid fibril from a lambda 1 light chain |
| 6z1o | AL amyloid fibril from a lambda 3 light chain in conformation A |
| 7nsl | AL amyloid fibril from a lambda 1 light chain |
| 8cpe | CryoEM structure of AL55 amyloid fibrils extracted from the kidney of an AL amyloidosis patient |
| 8r47 | AL amyloid fibril from the FOR010 light chain |
| 9eme | AL amyloid fibril from the FOR103 light chain |
| 9faa | Cryo-EM structure of cardiac collagen-associated amyloid AL59 |
| 9fab | Additional cryo-EM structure of cardiac amyloid AL59 - bent polymorph |
| 9fac | Additional cryo-EM structure of cardiac amyloid AL59 - mixed polymorph |

**Supplementary Table 3.**
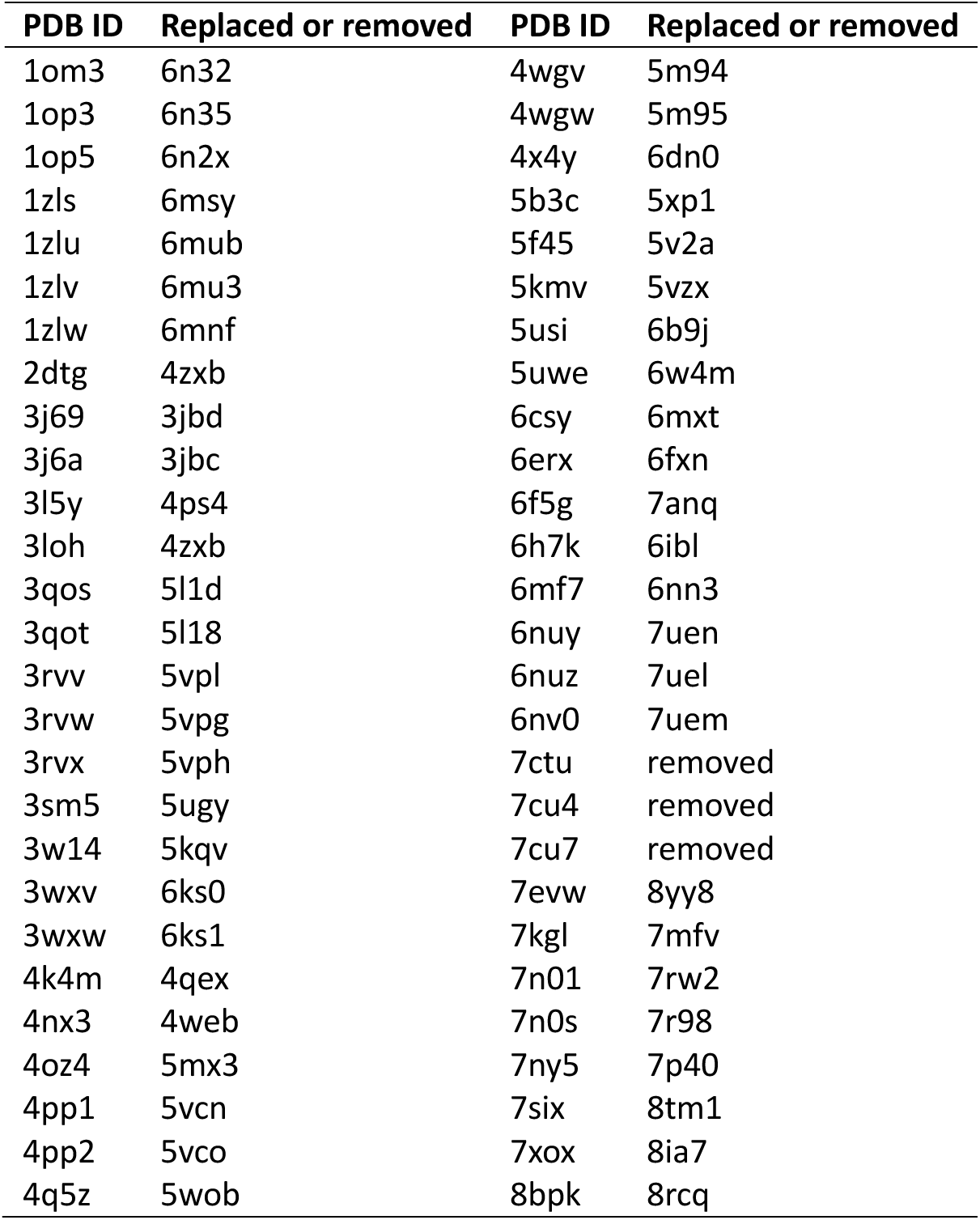
List of 54 PDB entries that have been replaced or removed.

**Supplementary Table 4.** List of 19 PDB entries containing multiple structural models.

| **PDB ID** | **Num of Antibody entries in SAAINT-DB** | **HC-LC pairings** | **Number of structural models in each assembly** | **Number of Antibody entries in SAbDab** |
| --- | --- | --- | --- | --- |
| 1g9e | 1 | A:N.A. | 20 | 20 |
| 1ieh | 1 | A:N.A. | 20 | 20 |
| 1maj | 1 | N.A.:A | 15 | 15 |
| 1mak | 1 | N.A.:A | 15 | 15 |
| 1vhp | 1 | A:N.A. | 20 | 20 |
| 2kh2 | 1 | B:N.A. | 77 | 77 |
| 2kqm | 2 | N.A.:A,<br>N.A.:B | 20 | 40 |
| 2kqn | 2 | N.A.:A,<br>N.A.:B | 20 | 40 |
| 2ltq | 2 | C:B, F:E | 20 | 40 |
| 2mkw | 1 | N.A.:A | 20 | 20 |
| 2mmx | 1 | N.A.:A | 20 | 20 |
| 2qtj | 4 | A:L, B:M,<br>C:N, D:O | 10 | 40 |
| 3chn | 4 | A:L, B:M,<br>C:O, D:N | 10 | 40 |
| 3cm9 | 4 | A:L, B:M,<br>C:N, D:O | 10 | 40 |
| 6iyn | 1 | A:N.A. | 20 | 20 |
| 6k2i | 1 | A:N.A. | 20 | 20 |
| 7eh3 | 1 | A:N.A. | 10 | 10 |
| 7v0v | 1 | A:N.A. | 20 | 20 |
| 8uek | 1 | A:N.A. | 20 | 20 |

**Supplementary Table 5.**
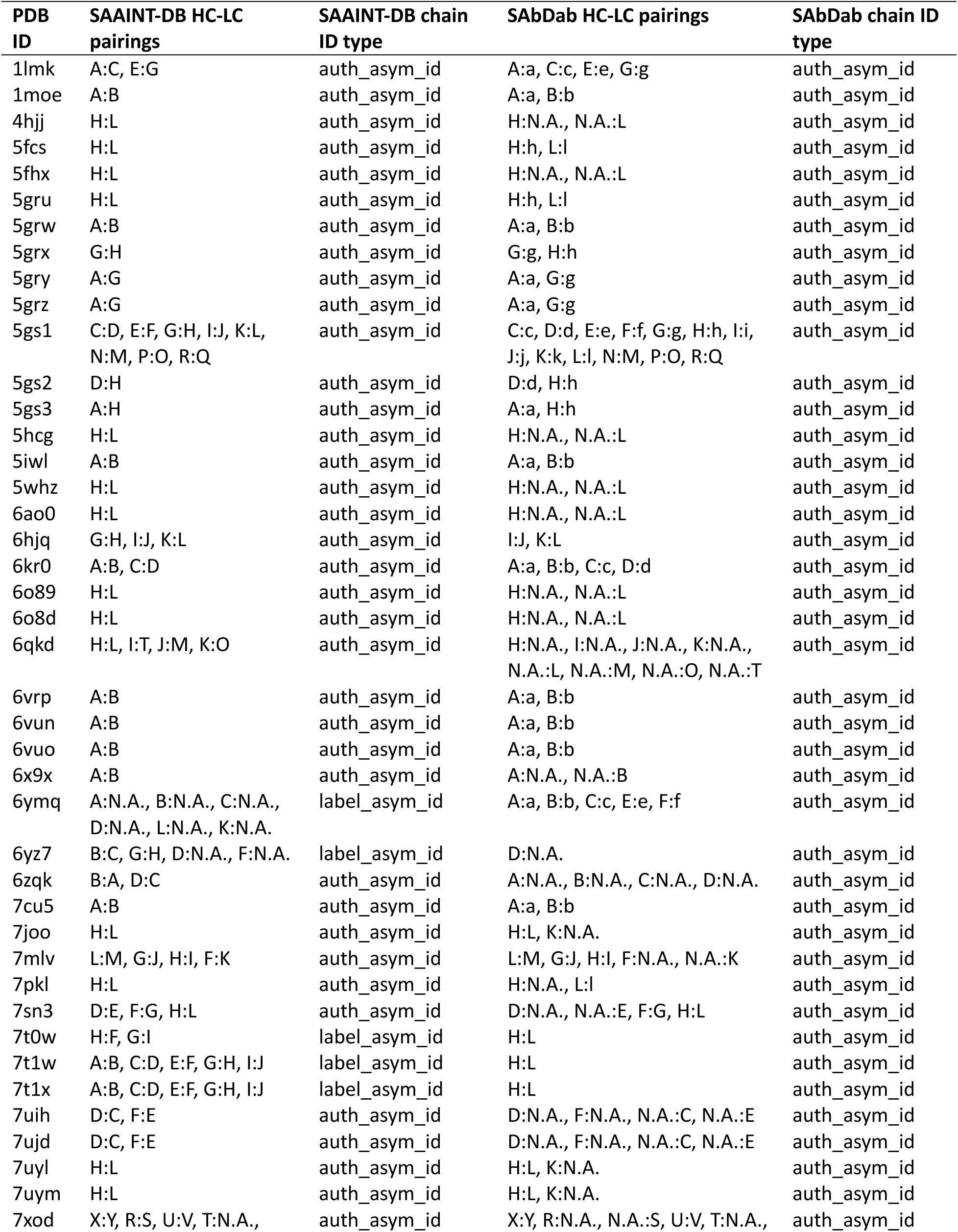
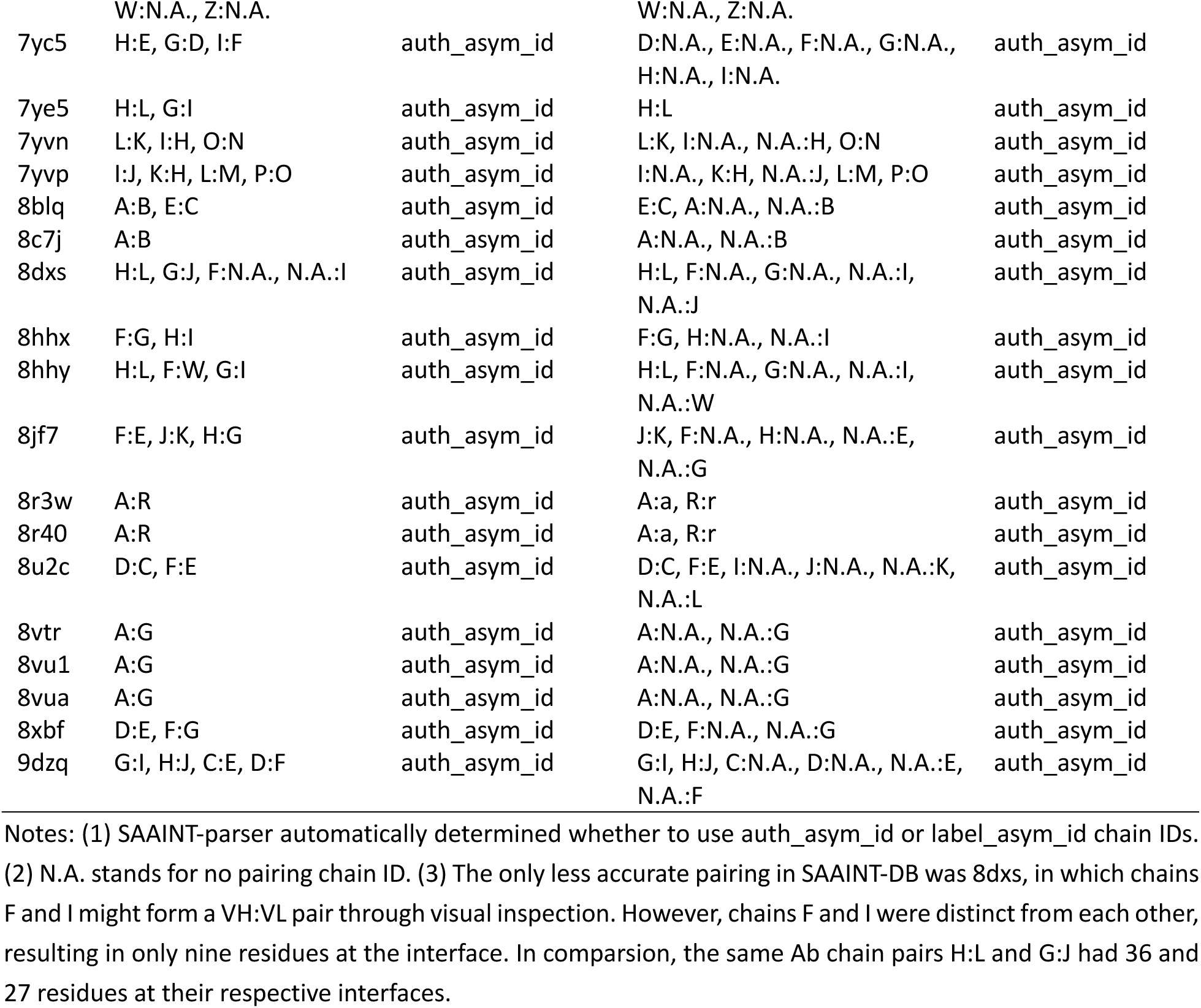
List of 60 PDB entries with incorrectly paired heavy and light chains in SAbDab, correctly paired in SAAINT-DB.

**Supplementary Table 6.**
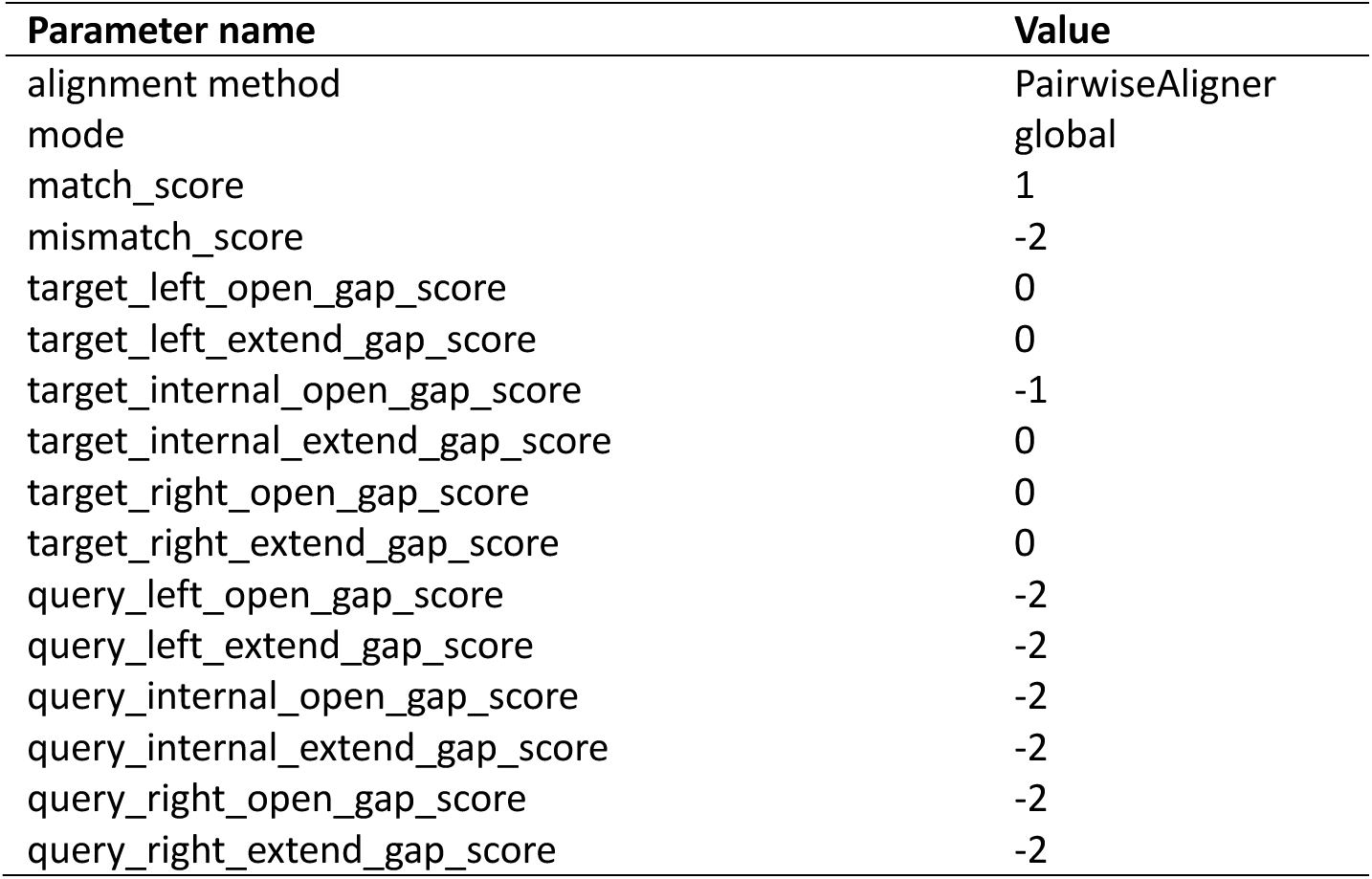
Customized parameters for pairwise sequence alignment.

| Parameter name | Value |
| --- | --- |
| alignment method | PairwiseAligner |
| mode | global |
| match_score | 1 |
| mismatch_score | -2 |
| target_left_open_gap_score | 0 |
| target_left_extend_gap_score | 0 |
| target_internal_open_gap_score | -1 |
| target_internal_extend_gap_score | 0 |
| target_right_open_gap_score | 0 |
| target_right_extend_gap_score | 0 |
| query_left_open_gap_score | -2 |
| query_left_extend_gap_score | -2 |
| query_internal_open_gap_score | -2 |
| query_internal_extend_gap_score | -2 |
| query_right_open_gap_score | -2 |
| query_right_extend_gap_score | -2 |

**Supplementary Table 7.**
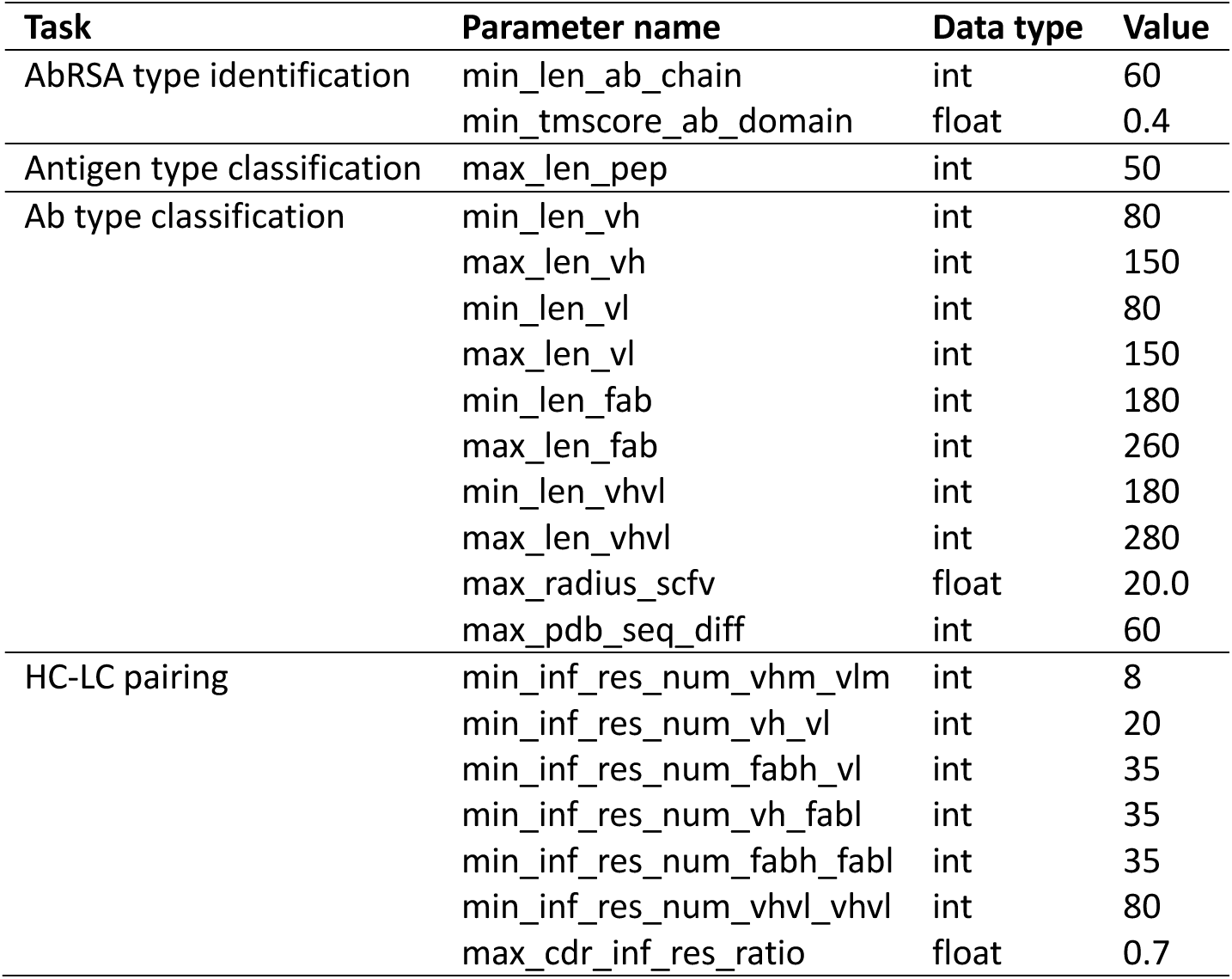
List of SAAINT-parser parameters.

**Supplementary Table 8.**
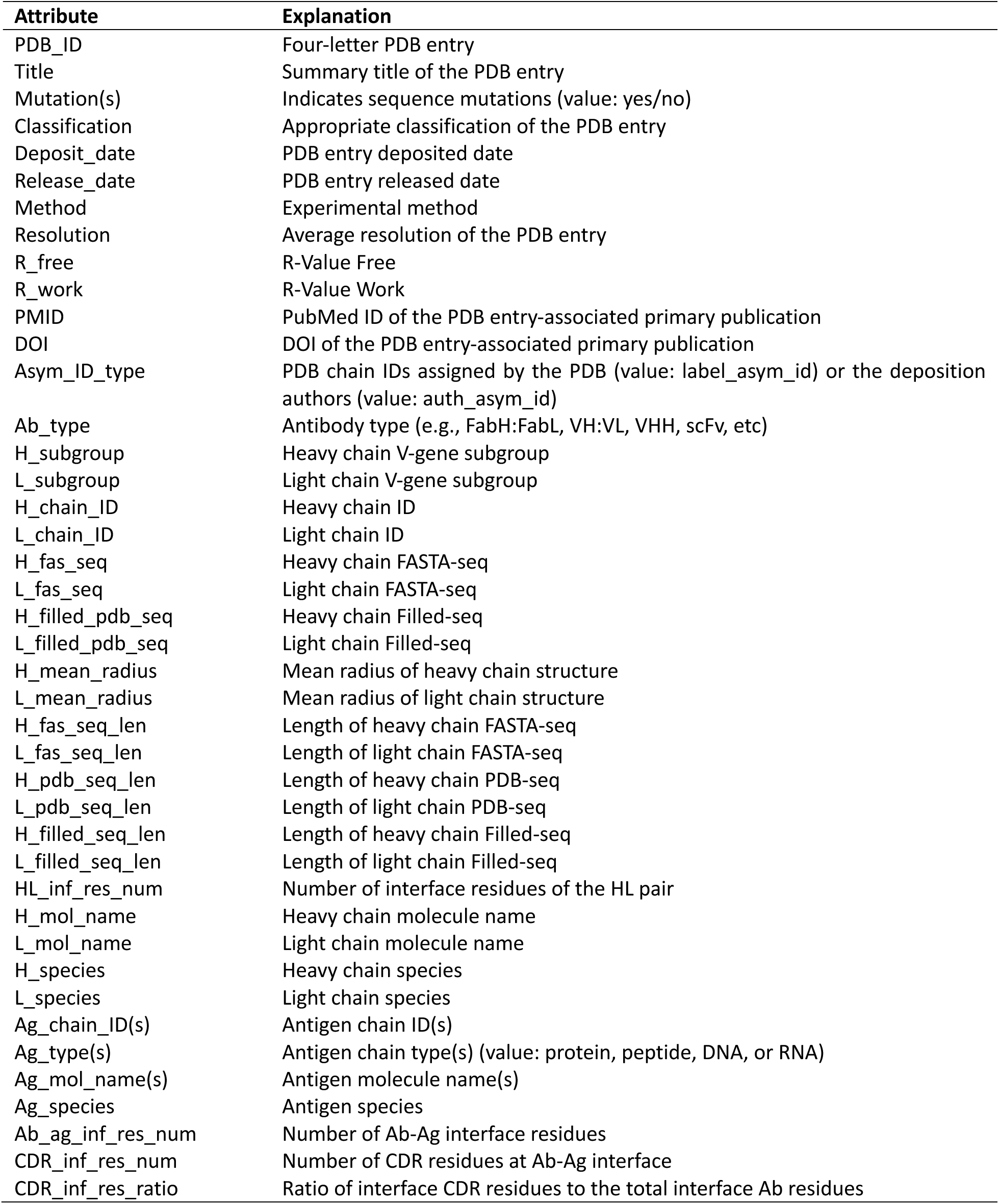
List of SAAINT-DB data entry attributes.

| **Attribute** | **Explanation** |
| --- | --- |
| PDB_ID | Four-letter PDB entry |
| Title | Summary title of the PDB entry |
| Mutation(s) | Indicates sequence mutations (value: yes/no) |
| Classification | Appropriate classification of the PDB entry |
| Deposit_date | PDB entry deposited date |
| Release_date | PDB entry released date |
| Method | Experimental method |
| Resolution | Average resolution of the PDB entry |
| R_free | R-Value Free |
| R_work | R-Value Work |
| PMID | PubMed ID of the PDB entry-associated primary publication |
| DOI | DOI of the PDB entry-associated primary publication |
| Asym_ID_type | PDB chain IDs assigned by the PDB (value: label_asym_id) or the deposition authors (value: auth_asym_id) |
| Ab_type | Antibody type (e.g., FabH:FabL, VH:VL, VHH, scFv, etc) |
| H_subgroup | Heavy chain V-gene subgroup |
| L_subgroup | Light chain V-gene subgroup |
| H_chain_ID | Heavy chain ID |
| L_chain_ID | Light chain ID |
| H_fas_seq | Heavy chain FASTA-seq |
| L_fas_seq | Light chain FASTA-seq |
| H_filled_pdb_seq | Heavy chain Filled-seq |
| L_filled_pdb_seq | Light chain Filled-seq |
| H_mean_radius | Mean radius of heavy chain structure |
| L_mean_radius | Mean radius of light chain structure |
| H_fas_seq_len | Length of heavy chain FASTA-seq |
| L_fas_seq_len | Length of light chain FASTA-seq |
| H_pdb_seq_len | Length of heavy chain PDB-seq |
| L_pdb_seq_len | Length of light chain PDB-seq |
| H_filled_seq_len | Length of heavy chain Filled-seq |
| L_filled_seq_len | Length of light chain Filled-seq |
| HL_inf_res_num | Number of interface residues of the HL pair |
| H_mol_name | Heavy chain molecule name |
| L_mol_name | Light chain molecule name |
| H_species | Heavy chain species |
| L_species | Light chain species |
| Ag_chain_ID(s) | Antigen chain ID(s) |
| Ag_type(s) | Antigen chain type(s) (value: protein, peptide, DNA, or RNA) |
| Ag_mol_name(s) | Antigen molecule name(s) |
| Ag_species | Antigen species |
| Ab_ag_inf_res_num | Number of Ab-Ag interface residues |
| CDR_inf_res_num | Number of CDR residues at Ab-Ag interface |
| CDR_inf_res_ratio | Ratio of interface CDR residues to the total interface Ab residues |

